# More Efficient Walking via Temporal and Spatial Energy Transfer in a Passive Biarticular Exosuit

**DOI:** 10.64898/2026.04.25.720622

**Authors:** Vahid Firouzi, Arjang Ahmadi, Ayoob Davoodi, Dennis Haufe, Andre Seyfarth, Gregory S. Sawicki, Maziar Ahmad Sharbafi

## Abstract

Evaluating bioinspired design principles in wearable assistive devices provides a unique opportunity to interrogate our understanding of the critical factors that enable agile, stable, and economical human movement. We introduce the BiArticular Thigh EXosuit (BATEX), a wearable device integrating two morphological features found in biological legged systems: biarticular muscles and elastic tissues. BATEX employs two biarticular springs spanning the hip and knee to emulate the human rectus femoris and hamstring muscles, creating beneficial synergy to enhance walking economy. This design enables two energy-shuffling mechanisms: temporal (spring-like storage/return at a joint) and spatial (strut-like transfer across joints). In walking experiments at **1.3 *m/s*** with N = 9 participants, a single compliant biarticular spring yielded a 7% metabolic cost reduction compared to walking without BATEX. Individually optimized configurations further improved metabolic reduction to 9%. BATEX morphology allowed users not only to off-load biological joint power (Assist) but also to increase total power (Augment). Across all exosuit configurations, the mechanical impact of the exosuit was reflected by a significant correlation between changes in users’ biarticular muscles’ activity and changes in net metabolic rate. In sum, compliant-biarticular exosuit architectures can concurrently assist and augment human lower-limb joint function, providing significant metabolic savings during walking.

## 1 Introduction

Biological systems feature embodied musculoskeletal structures integrated with neural control architectures enabling emergent agile, stable, and economical locomotion that engineered locomotor systems struggle to replicate [1]. Nature offers valuable blueprints for designing better robotic systems - especially for wearable robots that are intended to improve human locomotion performance by way of seamless human-robot interaction [2]. For example, it is possible to improve Human-Robot Interaction (HRI) by replicating the biological mechanisms that enable effective mechanical energy flow through the integrated neuromechanical structures within human limbs to produce smooth, steady locomotion [3]. Although non-biological design approaches (e.g.,[4, 5, 6]) have demonstrated advantages in specific contexts, aligning engineered systems more closely with biological principles could further enhance coordination between a human user and the wearable robot, ultimately accelerating the development of high-performance human augmentation with wearable technology for everyday use.

The integration of engineered and biological systems indeed presents significant challenges in developing wearable robots aimed at enhancing or restoring human mobility. This Challenge in HRI stems from the fundamental differences in structure and materials between these two systems, causing coordination gaps [7]. The large discrepancy between engineered and biological actuators in terms of both their intrinsic mechanical response and range of dynamic behavior has led researchers to prioritize the design of powered mechatronic systems with high-bandwidth torque/force control over unpowered bioinspired systems focusing on motion control synthesis [8, 9]. This approach, while effective, necessitates complex sensing, actuation, and control mechanisms to accurately replicate human locomotion. Conversely, the integration of human musculoskeletal properties into the design of assistive devices, such as exosuits, offers a promising alternative by fostering a synergistic human-exo interaction [10]. This method emphasizes “morphological computation” [11], harnessing the inherent “mechanical intelligence” embedded in biological systems [12] to simplify control strategies and enhance the efficiency and effectiveness of assistive devices [13]. Such designs not only support natural locomotion but also facilitate the validation of biological locomotion theories, underscoring the potential for passive systems to achieve widespread acceptance and use in daily life due to their simplicity and effectiveness for various user groups [14, 15, 16]. By focusing on muscle-like drives that incorporate elastic components and intelligent morphological designs, including multiarticular structures, wearable robots can provide significant, cost-effective solutions for human gait assistance [17].

The role of the biological actuation system is crucial in generating efficient, agile, and robust locomotion [17]. The continuous conversion of kinetic and potential energy in human gait is optimized by elastic, flexible, and multiarticular muscular structures [20]. The spring-like functionality of the leg [21] illustrates a sophisticated distribution of compliance across the musculoskeletal system [22]. The remarkable human leg design and the biarticular muscle arrangement facilitate this challenging motor control problem [22, 19, 23] while enabling functional stability and energy efficiency in legged locomotion [24, 25]. This architecture, coupled with nonlinear muscle dynamics, underscores a complex interplay of axial and rotational forces (Fig. 1a). essential for efficient locomotion [19]. Mechanical coupling between adjacent joints with biarticular muscles is also a biological solution for transferring energy between two joints [26, 27, 17] and has been previously used in legged robots [28, 29, 30, 19]. While some advancements in incorporating elastic elements in the design of passive exoskeletons for ankle [4] and hip joints [31, 6] for energy saving, the exploration into multiarticular designs remains limited [32, 33, 34, 35]. This gap underscores a significant opportunity for innovation in wearable technology, particularly in designs that mimic the biarticular muscle’s ability to provide cost-effective, simplified control mechanisms for human gait assistance [36, 37, 38]. Here, we fill this gap by introducing the BiArticular Thigh EXosuit (BATEX), an unpowered hip-knee exosuit that leverages the advantages of two morphological features found in biological legged systems: biarticular muscle arrangements and elastic tissues (Fig. 1).

**Fig. 1.**
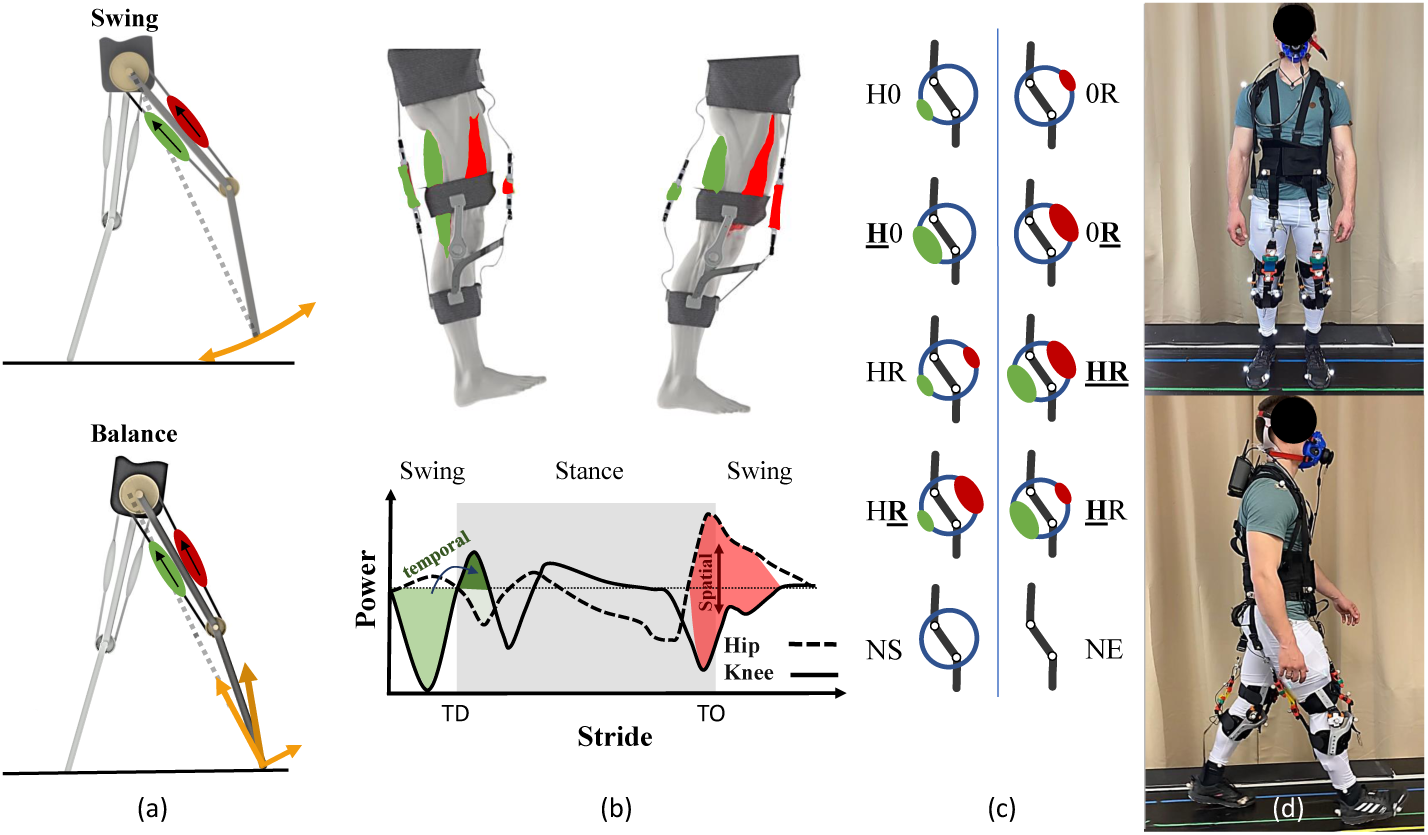
BATEX exosuit from conceptual design to experiment. **a)** Demonstrating the Contribution of the biarticular thigh muscles in walking with abstract models. The top figure conveys that biarticular thigh springs are sufficient to replicate human swing leg movement [18]. The bottom figure depicts the potential of the biarticular muscles to tune the ground reaction force (GRF) direction. Cross-talk of axial and tangential elements of the GRF could be minimized if the hip-to-knee lever arm ratio is set to two [19]. **b)** The mechanism of biarticular springs (mimicking hamstring (HAM, green) and rectus femoris (RF, red) muscles) in BATEX is demonstrated in the top figure. The bottom figure illustrates hip and knee joint power curves, highlighting the possible temporal and spatial energy transfer at these joints to be supported by the exosuit during the walking cycle. **c)** Different combinations of springs for HAM and RF in the BATEX. For unassisted cases, NS and NE represent No Spring and NO Exo cases, respectively. In the assisted configurations, the first and second letters refer to HAM (H) and RF (R), respectively. Three stiffness values are indicated by 0 for zero, regular font (H or R) for low and bold-underlined font (**<u>H</u>** or **<u>R</u>**) for high stiffness. **d)** The BATEX-assisted walking experimental setup includes metabolic cost measurement using the COSMED system, motion capture using infra-red high-speed Qualisys cameras, EMG, GRF from the instrumented treadmill, and Exo force measured by tensional force sensors. The individual pictured in this figure is an author of this study and has provided consent for publication.

The purpose of this study was to investigate the impact of BATEX on the metabolic cost of human walking. Drawing inspiration from the human musculoskeletal system, we placed BATEX’s biarticular springs (rubber bands) in parallel with the user’s rectus femoris (RF) and hamstring (HAM) muscles (Fig. 1b top). The design rationale centered around two key energy transfer mechanisms at the human hip and knee joints (Fig. 1b bottom): **(i) Temporal transfer with springs**, where elastic elements store energy during one phase of the gait and release it in another (e.g., in [16, 4, 39]) and **(ii) Spatial transfer with biarticular arrangements** enabling the energy absorption from one joint and transferring it to another (e.g., in [6, 38]). Notably, elasticity alone (e.g., in monoarticular joints) cannot enable spatial transfer, while biarticularity without elasticity (e.g., in [40]) does not allow for temporal energy storage and release. Therefore, combining elasticity and biarticularity can provide more effective assistance by enabling both temporal and spatial energy transfer. Based on our previous studies [37, 41, 42], stiff HAM and soft RF exosuit elements (e.g., Fig. 1c, **<u>H</u>**R) could provide optimal support for walking. The appropriate relationship between the leg segment lengths and moment-arm ratios of the two-joint muscles facilitates the decoupling of postural control from axial body support, crucial for balancing [17, 19] (Fig. 1a bottom). In addition, biarticular springs could simplify swing leg control by enabling morphological computation [18] (Fig. 1a top). Thus, we hypothesized that the biarticular springs of BATEX can significantly reduce the metabolic cost of walking without the need for an external power source by utilizing natural spatiotemporal energy flows between the hip and knee joint to reduce the effort. To test this hypothesis, we evaluated the response of nine healthy young adults walking at 1.3 m/s with the BATEX exosuit in nine unique biarticular spring configurations (Fig. 1c). Concerning the inter-subject variability, the different configurations allow us to identify the most effective spring arrangement for optimizing metabolic efficiency, with comparisons made to the no exo (NE) condition to assess the relative improvements across all tested conditions. Performance measures included net metabolic rate, lower limb joint mechanics, and lower limb muscle activity (Fig. 1d). In this process, we employed a discrete optimization procedure by cycling through all configurations to identify the most effective setup, a method inspired by Human-in-the-Loop optimization (HILO) [43] but adapted to optimize physical morphology considering the constraints of a passive system. Eventually, the verification experiments in this study validate the optimization process while addressing intersession and intra-subject variability—an aspect often overlooked in previous studies [31, 4, 5, 6, 36, 37, 38]. This approach opens up new possibilities for more reliable and personalized solutions in assistive robotics.

## 2 Methods

### 2.1 Participants

Nine healthy adults (AVG± SD; 9 male; age 27.9 ± 4.56 years; mass 73 ± 7.12 kg; height 1.8 ± 0.06 m) participated in this study. This sample size, which aligns with similar studies in the field [3, 4, 6, 44, 38], was determined based on prior research, pilot data, and practical constraints, ensuring sufficient variability to detect meaningful differences in metabolic cost and mechanical performance across the nine configurations. All participants voluntarily provided written informed consent before participation in this study. This study was approved by the Ethical Committee of the Technical University of Darmstadt and was carried out based on the guidelines of the Declaration of Helsinki.

### 2.2 BATEX hardware

The Biarticular Thigh Exosuit (BATEX) is designed to simultaneously assist the hip and knee joints by incorporating two springs per leg, as depicted in Fig. 1. One spring runs parallel to the human hamstring (HAM) muscle group, facilitating hip extension and knee flexion, while the other runs parallel to the rectus femoris (RF) muscle, supporting hip flexion and knee extension. The BATEX system weighs 2.2 *kg* and consists of rubber bands (acting as biarticular artificial muscles), textile components (shank, waist, and thigh braces), and articulated knee joints. The rubber bands are attached to the waist and shank braces via straps, forming the biarticular artificial musculature. A two-segment rigid linkage connects the shank brace to the thigh brace at the knee joint, stabilizing the biarticular spring’s endpoint on the shank brace and preventing undesirable movements. A 3D-printed knee lever is fixed to the shank brace to prevent direct contact and pressure from the RF artificial muscle on the patella, while also increasing the lever arm of the artificial RF for knee extension. Users can independently don the BATEX within five minutes after familiarization.

In this study, two sets of rubber bands with different stiffness values were selected: low stiffness (650*N/m*) and high stiffness (1100*N/m*). Preliminary experiments with two pilot subjects were conducted to determine the appropriate stiffness range. Participants’ oral feedback was used to select this range by progressively increasing the rubber bands’ stiffness during treadmill walking. The low stiffness was identified as the threshold where participants first perceived the exosuit’s assistance, while the high stiffness was determined as the upper limit before discomfort was reported. The realized stiffness at different joints depends on the varying moment arms and the individualized movement dynamics, as changes in spring length are a function of both the knee and hip joints. In Table. S1, we present the range of lever arms measured at different exosuit muscles on the hip and knee joints, based on kinematic data.

### 2.3 Protocol

Each participant completed two experimental sessions of *Main Experiment* and *Verification Experiment* on different days. Each of these experiments includes 10 trials of treadmill walking at 1.3*m/s*. Each trial consists of two minutes standing, eight minutes walking, and one minute standing, followed by three minutes allocated for spring adjustments. The sequence of 10 experimental conditions was randomized for each participant, with the no exo (NE) case placed at either the start or end of the experiment to accommodate the time required for exosuit fitting. In addition to the six-minute breaks between trials, we considered one 10-minute rest on a chair after the fifth trial. We did not consider the training phase for individual assistive scenarios. To prevent the springs from running slack, we adjusted their rest length so that they were slightly tensioned (not larger than 2*N*) when the participants were in a standing posture, with both knee and hip joints at straight configurations. Before each experiment, we conducted a short test (one minute) where the participant walked on the treadmill at the set speed, and we measured the force generated by the springs in real-time. We checked for substantial discrepancies between the expected and actual force patterns, and if necessary, we retuned the springs. Throughout the experiments, we continuously monitored the force patterns to detect any slack issues in real-time. Any significant slack issue was addressed, and the trial was repeated. Due to the rigid structure at the knee joint and the stable attachment points at the waist, the movement of the attachment point on the shank was minimal, further reducing the risk of slack.

#### Main Experiment

This experiment includes ten conditions: normal walking without wearing the exosuit and assisted walking while wearing the exosuit containing 9 combinations of three spring values (zero, low, and high) of two HAM and RF biarticular artificial muscles (see Fig. 1c). These conditions are abbreviated using two letters: NE (No Exo) and NS (No Spring) denote the unassisted and unsprung cases, respectively. For configurations involving at least one spring, ’0’ represents zero stiffness (spring removal), ’H’ as the first letter denotes the HAM muscle, and ’R’ as the second indicates the RF muscle. Standard typeface (H or R) represents low stiffness, while bold and underlined (**<u>H</u>** or **<u>R</u>**) denotes high stiffness, as shown in Figure 1.

#### Verification Experiment

In this experiment, we selected five walking conditions. In addition to the NE and NS cases, two configurations with fixed spring combinations were selected, which showed the highest average metabolic cost reduction in the Main Experiment with one spring (H0) and two springs (**<u>H</u>**R). The last configuration was the *Best* compositions of springs for each subject identified in the Main Experiment (unless it was H0 or **<u>H</u>**R). Each walking condition from the *Verification Experiment* was executed twice to calculate the average metabolic rate for each configuration.

### 2.4 Data collection

During the Main Experiment, we measured optical marker trajectories, ground reaction forces and moments, muscle activity, exosuit force, and metabolic energy consumption. In the verification experiment, just metabolic cost is measured. Motion capture, electromyography, and force data were time-synced using a triggering signal (voltage change) from the EMG system and recorded separately from the metabolic and device data. All data were segmented and normalized to 0–100% of the gait cycle.

We placed 23 reflective markers on the joints and bony landmarks to monitor the subject’s kinematics. A pair of additional markers were placed on the top and bottom of each rubber band to monitor the change in the rubber band’s length. Marker data were collected at 200 Hz with an optical motion capture system with 11 high-speed cameras (model OQUS300+ and OQUS310+, Qualisys, Sweden). Joint angles were calculated by tracking marker trajectories using a scaled model *Gait2354* in OpenSim software. Ground reaction forces were recorded at 1 kHz by the instrumented treadmill (ADAL-WR, HEF Tecmachine, Andrezieux Boutheon, France)) and at 100 Hz by the custom-built measurement adapter. All GRF trajectories were filtered using a zero-lag fourth-order low-pass Butterworth filter with 15 Hz cut-off frequency. We used the inverse dynamics tool in OpenSim software with a 6 Hz cut-off filter setting for the kinematics to calculate the joint torques. To measure the applied force (at 2 kHz), separate force sensors (KM10Z-200N, ME-System GMBH, Germany) were connected in series to rubber bands while attached to the shank brace on the other ends. All force trajectories were filtered using a zero-lag fourth-order low-pass Butterworth filter with 15 Hz cut-off frequency. The BATEX Kinetic sensory data was sent to a PC in real-time and monitored for adjusting springs’ rest lengths and stored via LabView software. Delivered torques generated by the exosuit during the assisted condition were calculated for each participant as the product of the force recorded by the load cells and the computed moment arms. Moment arms were defined as the perpendicular distance between the markers on the rubber band and the respective joint center.

The activity of 6 lower-limb muscles is collected at 2 kHz using wireless surface electromyography (EMG) sensors (Trigno Avanti, Delsys, Natick, MA, US). The 6 muscles recorded were: hamstring (HAM), rectus femoris(RF), vastus medialis (VAS), gluteus maximum (GLM), soleus (SOL), medial gastrocnemius (GAS). Electrodes were placed following guidelines in SENIAM (http://seniam.org/). EMG signals were bandpass filtered (fourth-order Butterworth, cut-off 20–450 Hz), rectified, and low-pass filtered (fourth-order Butterworth, cut-off 6 Hz) to obtain EMG linear envelope [45]. EMG signals were normalized by the average of corresponding EMG peaks recorded during the NE condition.

The metabolic cost was measured by an indirect calorimetry system (K5, Cosmed, Roma, Italy). Carbon dioxide and oxygen rates were averaged across the last 3 minutes of each walking condition and then used to calculate metabolic rate using the Brockway equation [46]. The net metabolic rate for each condition was obtained by subtracting the standing metabolic power from the walking metabolic power of each condition and then normalizing it by each participant’s body mass.

### 2.5 Analysis

#### Statistical analysis

To compare the results in different respects, we used a linear mixed effect model (LMM), where exosuit conditions were treated as categorical fixed effects and subjects were treated as random effects to account for subject-specific variability. In the following we elaborate on different aspects of our statistical analyses.

To compare the metabolic cost of each condition against the baseline configuration (NE or NS), we conducted pairwise comparisons using LMM. The model was specified as:

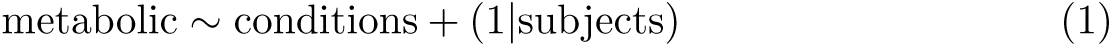

This model was implemented using the *fitlme* function in MATLAB, where the dependent variable (metabolic cost) was predicted by the conditions (categorical fixed effect) and random intercepts for subjects (random effect).

To correct for multiple comparisons, we applied adaptive False Discovery Rate (FDR) in the main experiment and standard FDR in the verification and averaged datasets. Adaptive FDR was chosen for the main experiment due to a low estimated proportion of true null hypotheses (*π*_0_ = 0.5), suggesting a potential power advantage. In contrast, standard FDR was used for verification and averaged analyses, where the benefits of adaptive FDR are less pronounced with *π*_0_ = 0 in fewer comparisons (*n* = 3).

To assess whether condition order influenced results in the main experiment, we used a linear mixed-effects model with condition order as a fixed effect and subjects as a random effect. This helped determine if order effects were significant while accounting for individual variability. The results indicate that there was no significant effect of experiment order on the outcomes (*p* = 0.968).

For analyzing the results of the Best condition, the same statistical method used for each configuration is applied to the Best condition. Since the statistical difference between the Best and NE or NS conditions in the main experiment may be artifactual due to selection bias, these cases are excluded from the analysis to avoid misleading information. For reporting the exosuit assistance effects for either fixed stiffness or personalized “Best” parameters, we consider three samples for each condition (two repetition from Verification and one from Main experiments) per participant. For each specific condition tried in both Main and Verification experiments, we used 27 data points (3 trials × 9 participants) and LMM was used for the pairwise comparison between unassisted and each of the assisted cases.

For statistical comparisons of Inverse Kinematic, Inverse Dynamic, and power curves across exosuit configurations relative to the NS condition, we performed pairwise comparisons at each time point using a linear mixed-effects model. Exosuit configuration was treated as a fixed effect, with subjects modeled as a random effect. This analysis produced a set of p-values across the gait cycle (100 time points × 8 comparisons). To control for Type I error, we applied adaptive FDR correction. Significant intervals were identified based on a predefined criterion: if a minimum number of two consecutive time points had p-values below 0.05, the interval was considered a significant deviation from the NS condition. This approach reduces the likelihood of isolated false positives and ensures that detected differences are sustained over time. Significant intervals are marked by horizontal lines corresponding to each exosuit configuration.

For the statistical analysis of EMG signals, we focused on phases where the muscles were actively engaged, specifically analyzing EMG peak values. Pairwise comparisons were conducted at these peaks using a linear mixed-effects model, with exosuit configuration as a fixed effect and subjects as a random effect. To control for Type I error in multiple comparisons, we applied an adaptive FDR correction.

#### Assistance vs. Augmentation

To explain the impact of different support mechanisms of each configuration of the BATEX exoskeleton, We defined the Assistance and Augmentation terms as follows:

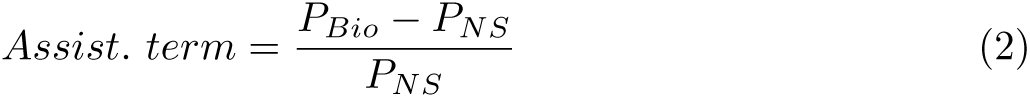

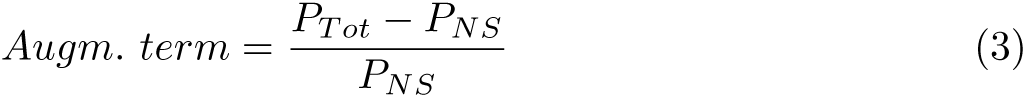

in which, *P_Bio_* and *P_Tot_* are the average of biological and total power of a certain joint at a determined time period for a specific configuration. The specified time periods are around touchdown (TD, swing-to-stance transition) or takeoff (TO, stance-to-swing transition), which are the critical periods of biarticular thigh spring contributions. More precisely, we defined the temporal energy transfer interval around TD, starting when the exosuit begins generating negative power at the knee joint (around 80% of the gait cycle) and ending when the subsequent positive power phase is completed (at 17% of the gait cycle). For spatial energy transfer analysis, we considered the interval from the point where the exosuit’s power at the hip and knee intersect (around TO, 50% of the gait cycle) until they cross again (at approximately 80% of the gait cycle). Subtracting the average biological power in the NS case in these periods (*P_NS_*) from *P_Bio_* and *P_T ot_* and normalizing it to *P_NS_* introduce a measure to evaluate how much reduction in biological power or increase in total power can be achieved by each configuration. Based on this definition, negative values for assistance and positive values for augmentation are desirable, as they correspond to reductions in biological power and increases in total power, respectively.

#### EMG-Metobolic analysis

To examine the link between muscle activity variations and metabolic cost due to BATEX support, a subsequent linear regression analysis was conducted. In this method, known as forward selection [47], muscles were sequentially incorporated based on their contribution to the highest increase in adjusted-*R*^2^ for the model’s accuracy. This selection process was carried out until the regression model included all six monitored muscles, as excluding any of them could miss important contributions to the overall metabolic expenditure. While allowing negative regression coefficients could improve correlation, the model was constrained to ensure positive coefficients (except for the bias term), to meet the physiological expectation that increased muscle activation raises metabolic cost [48, 49, 50]. After including all muscles, the adjusted-*R*^2^ was used to assess the exo’s effect on each muscle and understand BATEX’s functionality in reducing metabolic cost. This second refinement with adjusted-*R*^2^ ensures that the inclusion of each muscle adds meaningful explanatory power to the model without artificially inflating the fit. The participant-average fit equation, *R*^2^, and p-value were calculated using fitted data on changes in muscle activity versus changes in metabolic cost.

## 3 Results

To assess the biomechanical and energetic impact of the BATEX, we analyzed how different configurations and stiffness levels of biarticular springs affected users’ metabolic cost, joint power, kinematics and dynamics, as well as muscle activity during treadmill walking.

### 3.1 Metabolic cost

The BATEX reduced the metabolic cost of walking for a subset of the biarticular spring configurations. For all participants, the *Main Experiment* revealed at least one combination of BATEX HAM and RF springs that resulted in reduced metabolic cost of walking compared to no exo (NE) (Fig. 2, left), and this was confirmed in a follow-up *Verification Experiment* (Fig. 2, mid).

**Fig. 2.**
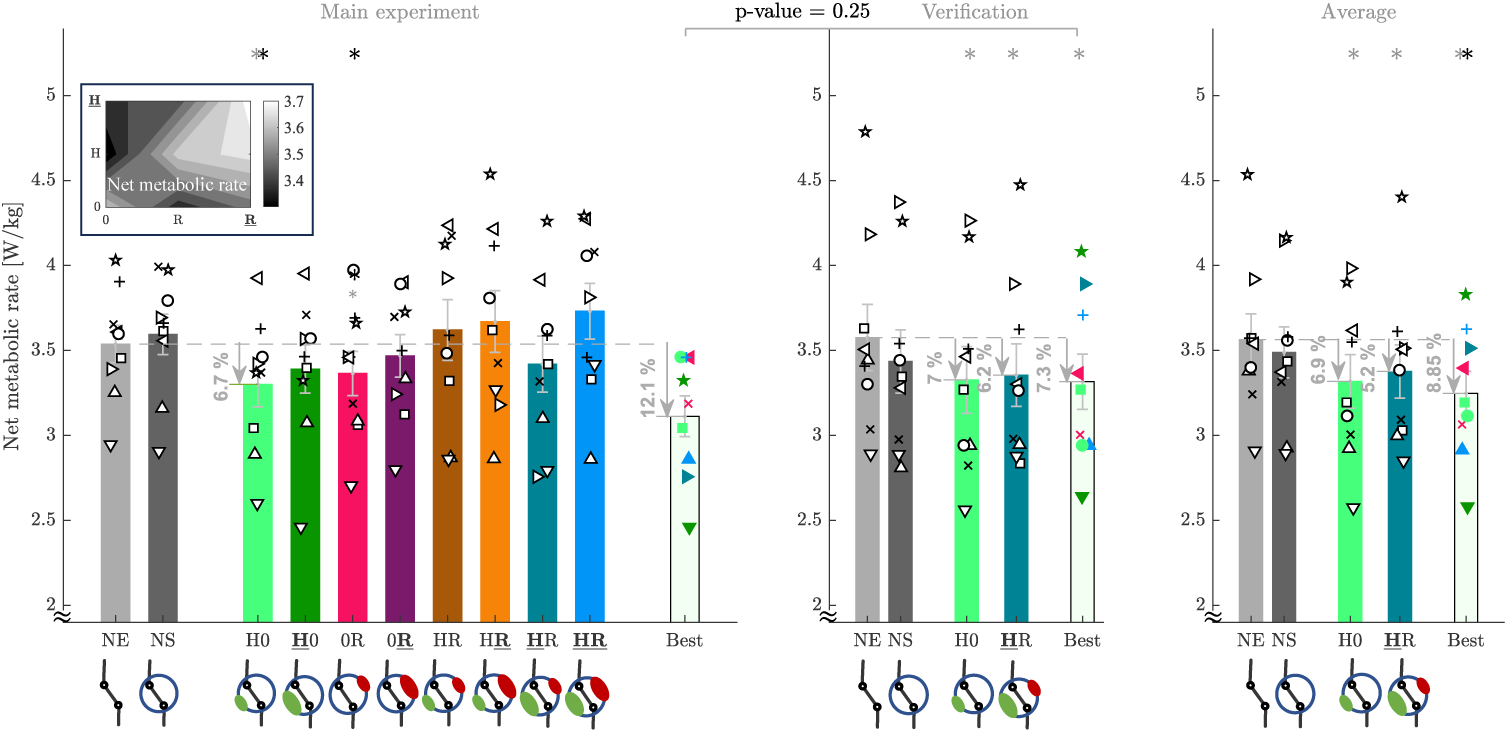
Human metabolic rate changes with assistance for waking at 1.3 m/s. The net metabolic rate is normalized with the body mass, and the values for quiet standing are subtracted in both experiments. Bar graphs show the average and variance among all nine participants, while the outcomes of each configuration are shown by a specific color. The metabolic rate for each participant is indicated by specific white markers at different conditions. For each participant, the *Best* configuration is the one with minimum metabolic consumption for that participant in the *Main Experiment*. The colors of the markers for the Best condition denote which configuration results in the minimum metabolic rate for that specific participant. **Left)** The *Main Experiment* with all configurations **Middle)** The *Verification Experiment* with the two times repetitions of five different configurations: NE (no exo), NS (No spring), H0 (low HAM stiffness without RF), **H**R (High HAM stiffness with low RF stiffness) and also the *Best* condition which minimizes the metabolic rate for each participant in the *Main Experiment*. The two fixed configurations of H0 and **H**R are selected as configurations with the lowest average metabolic consumption (in the *Main Experiment*) with one and two springs, respectively. **Right)** The average of the main experiment (one trial per condition) and verification experiments (two trials per condition). The horizontal dashed line indicates the average value for NE in each experiment, and ^∗^ shows *p <* 0.05.

In the Main experiment, across all participants, the H0 (low stiffness HAM) configuration (3.3 ± 0.13 *W/kg*) significantly reduced metabolic cost by 6.7 ± 2.57% (*p* = 0.033) compared to no exo (NE) (3.53 ± 0.11 *W/kg*). The low stiffness RF spring (0R) did not provide a significant metabolic benefit compared to NE, but it significantly reduced user effort by 6.24 ± 2.6% (*p* = 0.048) with respect to no springs (NS) (3.59 ± 0.12 *W/kg*). We selected the H0 and **<u>H</u>**R configurations as the ones with the lowest metabolic cost, respectively, among the single and two-spring configurations. Additionally, the individualized *Best* condition (with minimum metabolic rate) was determined for each participant. These five conditions (NE, NS, H0, HR, and Best) were further tested in the Verification experiment. The follow-up Verification experiment confirmed the results from the Main experiment (e.g., a 7.0 ± 2.2% reduction in the H0 vs. NE condition, *p* = 0.002, 3.3 ± 0.19 *W/kg* Fig. 2, right). In this experiment, **<u>H</u>**R also showed significant metabolic reduction of 6.2 ± 2.2%, compared to NE (*p* = 0.01, 3.33 ± 0.19 *W/kg*).

To report the final effects of assistance, both experiments were considered together, using the average of three samples per condition (two repetitions from the Verification and one from the Main experiment) per participant. The average reductions are shown for the five conditions in Fig. 2, right. All three assisted scenarios showed a significant reduction in metabolic cost compared to the NE condition: H0 (6.9 ± 2.1% reduction, *p* = 0.007, 3.3±0.16 *W/kg*), **<u>H</u>**R (5.2±1.9% reduction, *p* = 0.03, 3.35±0.15 *W/kg*), and the Best condition (8.9 ± 1.75% reduction, *p <* 0.001, 3.25 ± 0.13 *W/kg*) including different configurations across participants. No significant difference was found between the Best condition results from the Main and Verification experiments (*p* = 0.25).

### 3.2 BATEX contributions to lower-limb joint power

To analyze the effects of biarticular springs on kinematic and dynamic behavior, as well as energy management, we will hereafter compare different configurations with NS. BATEX spring configurations that provided metabolic benefit (H0, 0R and **<u>H</u>**R) demonstrated both temporal and spatial patterns of mechanical energy flow to assist and augment users’ hip and knee joint mechanical power compared to walking without exosuit springs (NS) (Fig. 3). In BATEX configurations with a biarticular hamstring spring (H0 (green), **<u>H</u>**R (blue)), the exosuit generated peak negative mechanical power at the knee joint that approached -0.4 W/kg and significantly reduced both biological (H0, 12.6%, *p <* 0.05 and **<u>H</u>**R, 27.3%, *p <* 0.05) power, while the total (H0, knee joint power does not significantly change in this terminal swing phase (Fig. 3, top left). A portion of the stored energy in the BATEX hamstring springs was subsequently returned, significantly reducing peak positive biological knee joint power (e.g., **<u>H</u>**R, 37.4%, *p <* 0.05) in the early stance phase. In general, the temporal transfer of stored BATEX spring energy from terminal swing to early stance provided assistance to users at knee joint (i.e., reduced biological power compared to walking with no springs (NS (black))). The assistance-augmentation behavior, characterized by changes in biological and total power, is illustrated in the bottom left of Fig. 3.

**Fig. 3.**
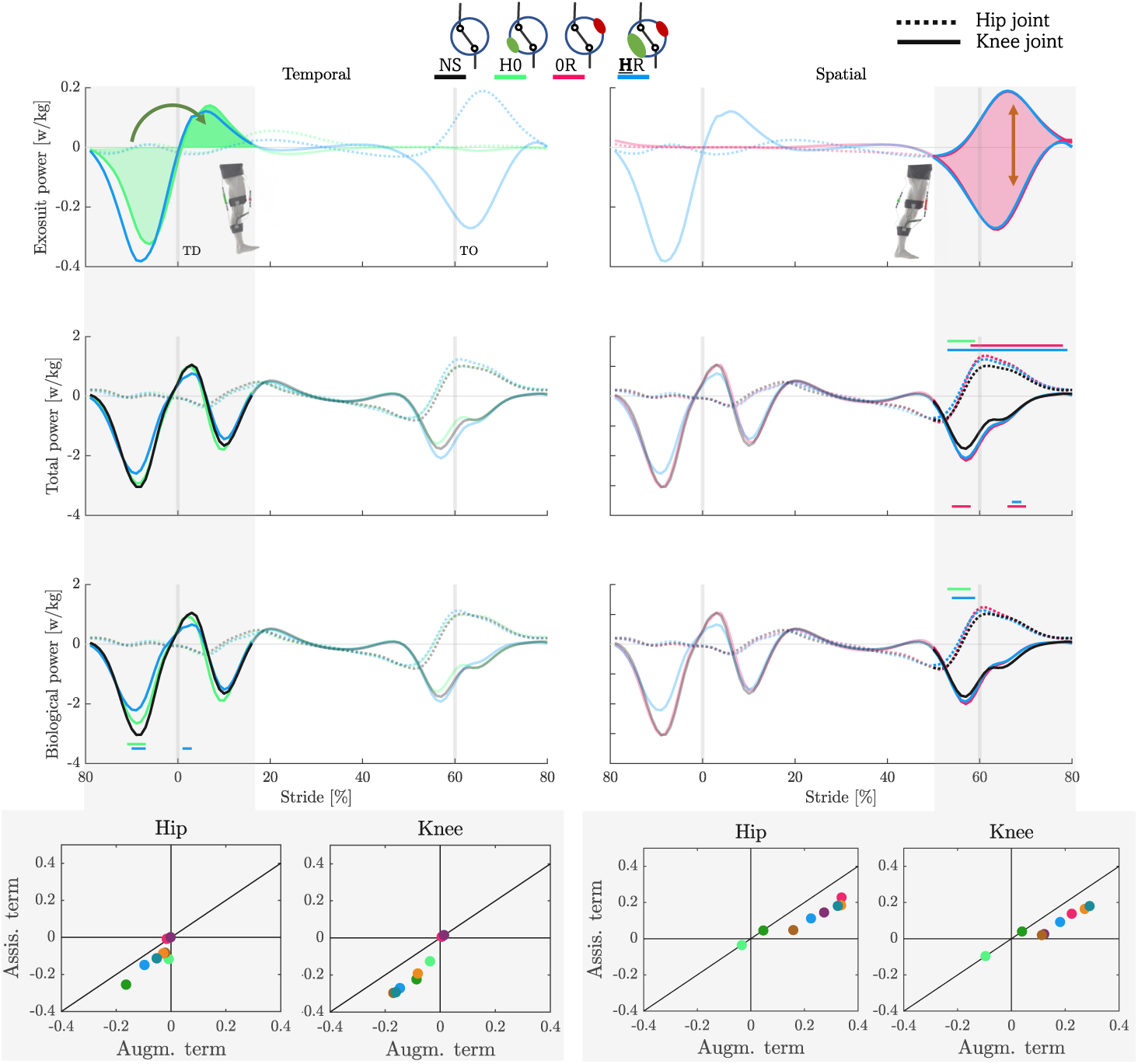
BATEX assistance and augmentation effects on hip and knee joints. The first three rows show the exosuit, biological, and total (Exo + human) power in one stride (average across participants), comparing H0, 0R, and **<u>H</u>**R (selected based on metabolic reductions) with the NS case. Vertical grey lines indicate touchdown (TD) and takeoff (TO) moments. Shaded grey regions highlight temporal and spatial energy transfer, respectively, in the left and right columns. Statistically significant differences (*p <* 0.05) between the NS condition and other exosuit configurations are marked by horizontal lines above (for the hip) and below (for the knee) joint curves. *Assist. term* (Eq. 2) and *Augm. term* (Eq. 3) are shown for the highlighted time periods (average of all participants). Each circle represents a specific configuration, color-coded as in Fig. 2. Positive *Augm. term* and negative *Assist. term* indicate augmentation and assistance, respectively. Circles below the unit slope line indicate exosuit and biological power alignment.

In BATEX configurations with a biarticular rectus femoris spring (0R (red), **<u>H</u>**R (blue)), the exosuit simultaneously generated peak positive mechanical power at the hip joint (∼ 0.2 *W/kg*) and peak negative mechanical power at the knee joint (∼ 0.3 *W/kg*) (Fig. 3, top right). BATEX mechanical energy transfer from the knee joint to the hip joint tended to increase both negative total knee joint power (e.g., in 0H, 22.1%, *p <* 0.05) and positive total hip joint power by (0H, 34.8%, *p <* 0.05 and **<u>H</u>**R, 23.3%, *p <* 0.05) in the stance to swing transition (50% − 70% stride). Its effects on increasing negative biological knee joint peak power and positive biological hip joint peak power during the same period are not significant. Overall, spatial energy transfer via BATEX biarticular rectus femoris springs amplified both total knee and hip joint powers compared to walking with no springs (NS (black), in Fig. 3, middle and bottom right).

### 3.3 BATEX-induced neuromechanical response at hip and knee joints

BATEX spring configurations that provided metabolic benefit (H0, 0R, **<u>H</u>**R) modified users’ biological joint moments, muscle activity and movement patterns compared to walking without exosuit springs (NS) (Fig. 4). In BATEX configurations with biarticular hamstrings (H0 (green), **<u>H</u>**R (blue)), the exosuit produced hip extension (Fig. 4, left top) and knee flexion torque (Fig. 4, right top) from mid-swing (80% stride) through mid-stance with peak values reaching 0.05 *Nm/kg*. BATEX hamstring springs can significantly reduce the biological knee flexion moment in late swing (Fig. 4, right second-row), e.g., **<u>H</u>**R with 30% reduction (*p <* 0.05) (see other conditions in Fig. S5). The effects of BATEX knee flexion/hip extension torque on increasing users’ hamstring muscle activity are not significant during early stance (Fig. 4, right third-row), while they significantly increase knee joint flexion (*p <* 0.05) during the whole stance phase (Fig. 4, right bottom).

**Fig. 4.**
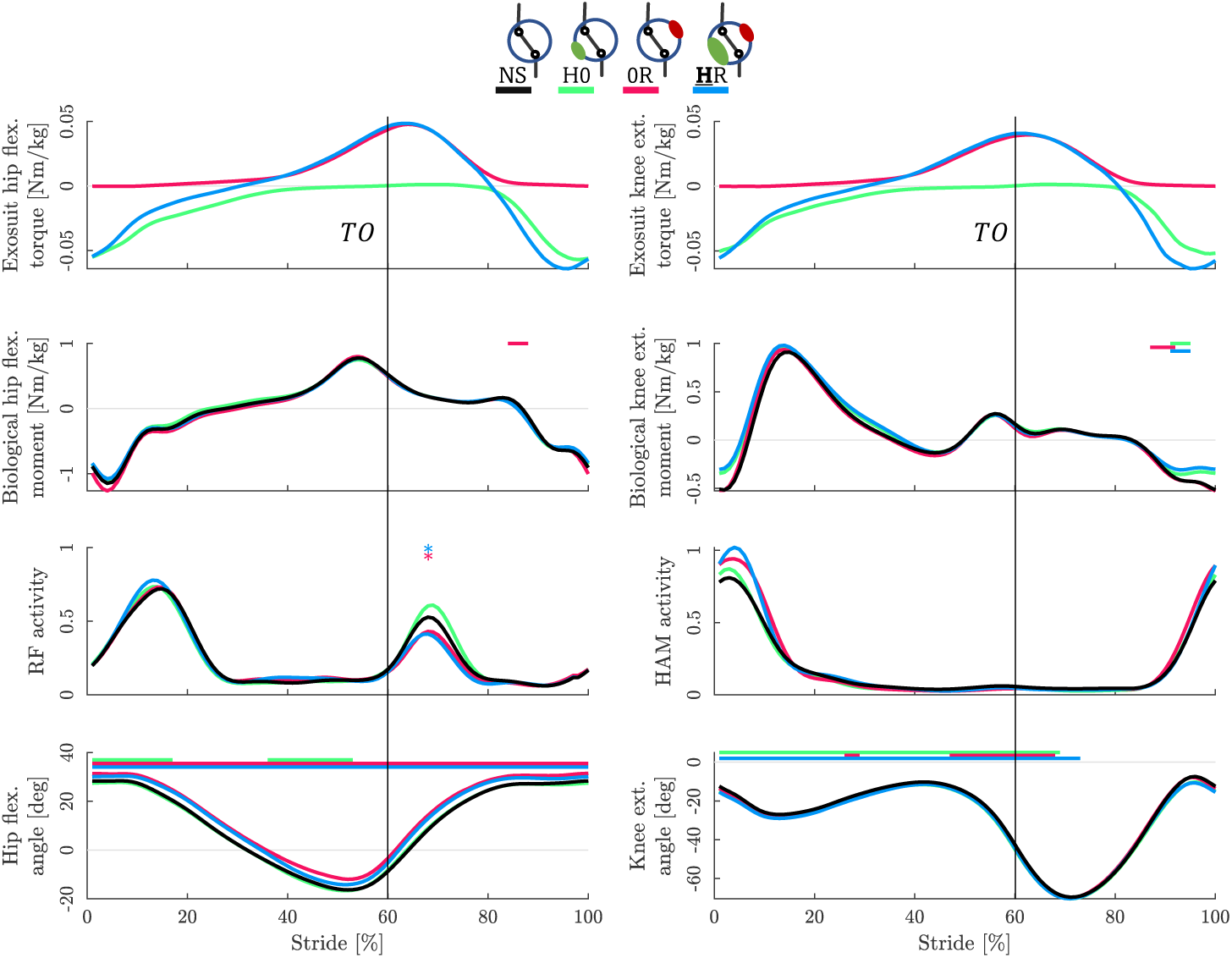
Mechanics and muscle activity. Time series of average values of exosuit and biological torque at hip and knee joints, biarticular thigh muscle activations, and joint angles of hip and knee joints in one stride for 0R, H0, and **H**L configurations (selected based on metabolic reductions), compared to NS (average of all participants). Statistically significant differences (*p <* 0.05) between the NS condition and other exosuit configurations are marked with horizontal lines above each figure. For EMG analysis, we focused on EMG peaks, with significance indicated by asterisks (*). The stride starts with a touchdown of one leg (0%), and the vertical black line indicates the takeoff of the same leg (60%). Torques at hip and knee joints (first and second rows) are normalized to body mass. Biological torques are calculated by subtraction of the exo torque from the total torque (found by inverse dynamics). Muscle activity in the rectus-femoris (RF) and hamstring (HAM) are selected as the main two muscles to be supported by the BATEX. Kinematic variations at the hip and knee joint angles are demonstrated as the joints with direct interactions with the exosuit.

With exosuit biarticular rectus femoris springs (0R (red), **<u>H</u>**R (blue)), the exosuit produced hip flexion (Fig. 4, left top) and knee extension torque (Fig. 4, right top) from mid-stance (40% stride) through mid-swing (80% stride) with peak values reaching 0.05 *Nm/kg*. BATEX rectus femoris springs had little effect on biological moments at either the hip (Fig. 4, left middle) or the knee (Fig. 4, right middle), but instead acted to increase the total moments at each joint in swing phase (see Fig. S6). With biarticular springs delivering BATEX knee extension/hip flexion torque, users significantly reduced rectus femoris muscle activity in early swing (*p <* 0.05) (Fig. 4, left third-row), and significantly increased hip joint flexion angle throughout the entire gait cycle (*p <* 0.05)(Fig. 4, left bottom).

### 3.4 BATEX user Δ lower-limb muscle activity vs. Δ net metabolic rate

The changes in BATEX users’ lower-limb muscle activity reflected changes in net metabolic rate during walking with the device across all spring configurations. We found a significant relationship between changes in cumulative lower-limb muscle activity and changes in net metabolic rate (*p* = 0.01; *R*^2^ = 48.1%) – users that reduced lower-limb muscle activity the most with BATEX assistance demonstrated the largest metabolic benefits (Fig. 5, right – lower left quadrant). As seen in the regression, the most influential muscles were the rectus femoris (RF) and hamstrings (HAM), together having *R*^2^ = 44.4%, confirming their key role in the exosuit’s mechanical assistance. Adding other muscles slightly increases the correlation by gastrocnemius (GAS, 2%) and gluteus maximus (GLM), soleus (SOL), and vastus (VAS), each contributing less than 1% (Fig. 5, left). Including all six muscles in the regression model captured the full effect of muscle activations on metabolic cost, as expected, while adding SOL and VAS reduced the adjusted-*R*^2^.

**Fig. 5.**
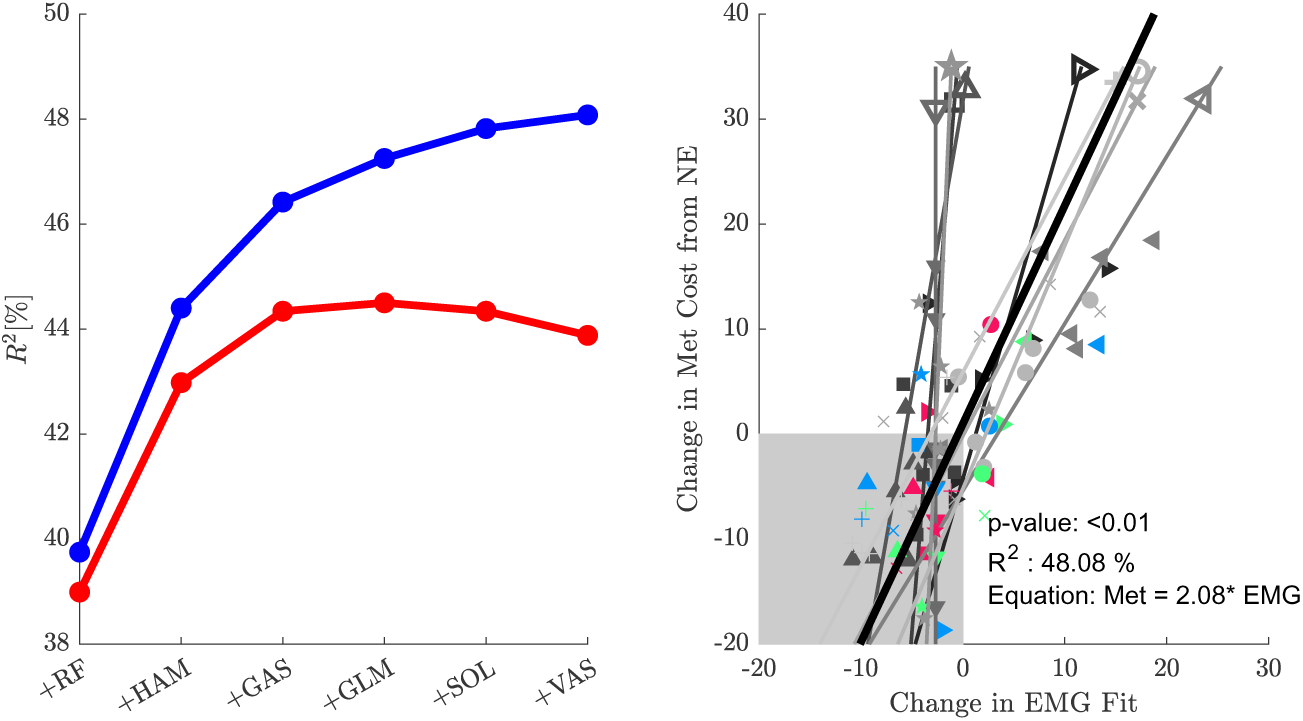
Association between changes in users’ lower-limb muscle activity and metabolic cost across 10 different assistance conditions. Left) Participant average *R*^2^ (blue) and adjusted *R*^2^ (red) values, generated through iterative regression to link variations in muscle activity per gait cycle (in %) and alterations in overall metabolic rate (in %), benchmarked against the condition without the Exosuit (NE). On the x-axis, the muscles included in the model are cumulative from left to right so that each muscle’s plotted *R*^2^ point corresponds with a model that incorporates all previously listed muscles. **Right)** Metabolic changes vs. EMG changes and linear approximation of their relation. Different trials are illustrated with markers while the colored ones highlight the three selected conditions (H0, 0R and **<u>H</u>**R), color-coded as Fig. 3. Individual participant linear regression outcomes (depicted with grey lines) and the collective average across all subjects (shown with black lines) are displayed. Grey-shaded squares underscore instances where metabolic rates decreased in comparison to the NE case.

## 4 Discussion

The overarching objective of this study was to demonstrate the ability of ’morphological computation’ to assist human locomotion. More specifically, we used the passive biarticular thigh exosuit (BATEX) to identify morphological configurations that, through mechanical coupling and interaction, reduce the metabolic cost of walking — without requiring external computation or control effort. Our hypothesis was that by using biarticular springs to optimize *temporal* and *spatial* energy transfer between the hip and knee joints, an unpowered exosuit could reduce the metabolic energy required for human walking.

### 4.1 Cost-less assistance

Our findings revealed that BATEX could leverage the principle of temporal and spatial energy transfer to reduce the metabolic cost of walking (Fig. 1b). Human users can use a wearable device to save metabolic energy - even if the device has no explicit energy source (i.e., no net work generating motors). For example, mechanical elements in parallel with human joints can reduce user joint moments/muscle forces and save musculotendon negative work by (i) permanently extracting energy with dampers [51, 39] or (ii) temporarily storing energy in passive compliant elements and then returning it later during a locomotion cycle (Fig. 1b, green) to also save positive work [26, 4], and (iii) transferring power from one joint to another (reducing internal work, Fig. 1b, red) [6, 44]. Indeed, BATEX could significantly reduce metabolic expenditure using a fixed, across-participant configuration (Fig. 2 right, H0 and **<u>H</u>**R) as well as by selecting the per-participant ‘Best’ configuration (Fig. 2, Best).

### 4.2 Temporal energy transfer with HAM

The HAM spring mainly employs the temporal energy transfer by storing joint power in one phase and returning it in the following phase. This energy transfer mechanism (shown by green shaded region in Fig. 3) involves a time shift, where, for example, energy is absorbed by the HAM spring during knee extension in the swing phase and returned in the early stance, thereby reducing the metabolic cost. As the hip angle is almost constant in this period (Fig. 4 bottom left), the HAM works similarly to a monoarticular knee flexor muscle. Instead of dissipating energy at the knee joint (e.g., using a damper [52, 53]), BATEX’s elastic behavior restores negative power to support knee flexion around swing to stance transition. Biological joint power curves indicate that with HAM spring (in **<u>H</u>**R and H0), knee negative power in the late swing phase and the positive biological power in the early stance phase will reduce. The power reduction is consistent with the significantly reduced knee torque at swing-to-stance transition (e.g., with **<u>H</u>**R, shown in Fig. 4). This temporal transfer of energy could also be beneficial for a new type of exosuits designed to assist prosthetic devices (e.g., ankle and knee prostheses, [54]) supported by a biarticular thigh exo like BATEX.

### 4.3 Spatial energy transfer with RF

*Spatial energy transfer* is another potential mechanism humans could use to leverage an unpowered device and reduce the metabolic cost of walking. In this case, the device would transfer user energy from one joint to another joint, reducing the demands for internal work by providing a cost-less path for energy transfer [6] (Fig. 1b, red). Spatial energy transfer does not necessarily require an elastic mechanism, as seen in the nonelastic gastrocnemius-like string in the Jumping Jack model [40], where it enhances jump height by aligning the ground reaction force more vertically.

The RF spring mainly contributes to spatial energy transfer between two joints in the late stance and early swing, as seen in the exosuit power in Fig. 3. This exo contribution (e.g., in 0R and **<u>H</u>**R) increases the total joint power, resulting in a higher hip flexion angular velocity around take-off (see Fig. S10). The higher total power in the early to mid swing phase, which continues the augmentation capability of the exo via spatial energy transfer, aligns with the significant increase in total torque shown in Fig. S6. A comparable increase can also be observed in the total extension torque of the knee joint. Consequently, the second peak of RF muscle activation will be reduced (Fig. 4).

### 4.4 Assistance vs. Augmentation

To better understand the exo contribution to joint power management, we demonstrated the changes in total power resulting from BATEX contribution and its relation to biological power changes in separate phases. The *Assistance vs. Augmentation* graphs (Fig. 3, bottom left) highlight the assistance capability of the exo (defined as *Assist. term* in Eq. 2 of [55]) in the late swing and early stance. Thus, the temporal energy transfer supports Swing LSFs with breaking knee extension in late swing as can be seen by a significant reduction in biological knee power (Fig. 3) and knee torque (Fig. 4). Further, Stance LSF is supported by assisting knee flexion in the early stance phase as observed in a significant reduction of biological knee power in **<u>H</u>**R and **<u>H</u>**0 (Fig. S2).

The *Assistance vs. Augmentation* graphs (Fig. 3 bottom right) further explain the RF spring effects on power management (see Eq. 3 in [55] for *Augm. term*). The power generated by the exo with the RF spring leads to an additional surplus of biological power. This is demonstrated by the circles in the first quadrant beneath the diagonal line with slope one. Analyzing torque profiles (Fig. 4) helps to better understand the observed augmentation functionality. While the hip torques produced by RF springs are consistent with biological data, the positive hip torque peak occurs later than the biological one. This delay could explain the reduced RF muscle peak activation during the early swing phase (Fig. 4). However, the RF spring does not decrease hip flexion torque in the late stance and early swing phases. Instead, with lower biological RF muscle effort, the increased total hip torque (shown in Fig. S6) will accelerate the leg in the early swing phase S10). Nevertheless, the *augmentation capability* provided by the RF spring results in a significant increase in hip angular velocity peak, followed by a significant reduction of hip protraction speed, later in the swing phase (see Fig. S10). This latter effect could be a compensation to avoid increasing walking speed, given that the treadmill speed is fixed. Consequently, no significant differences were observed in gait phase timings (stance, swing, double-support phases) or step length. Further advantages of the BATEX augmentation capability, such as increasing walking speed, should be explored in future research, in addition to the achieved benefits like energy savings and reduction in RF muscle activation.

### 4.5 Generalizable vs. Individualized Tuning of Hip-Knee Exosuits

Our experiments support a growing number of studies [37, 36, 38, 44] and verify that unpowered wearable devices (e.g., exoskeletons or exosuits) attached at proximal joints (e.g., hip, knee) can reduce the metabolic cost of walking by about 9%, which is more than twice the reported assistance with passive monoarticular exoskeletons at the hip [56] and about three times the reduction reported for knee assistance with passive devices [52]. We expect future exosuit users will represent the natural population and thus will come in all shapes, sizes and physiological make-ups. Despite this potential challenge to a ‘one-size-fits all’ solution, our exhaustive examination of the multifactorial design space covering the full range of compliant, biarticular flexion-extension morphologies (Fig. 1c) singled out a core morphological feature that is key for *Best* performance *across* individual users. That is, in both our *“Main”* (Fig. 2, left) and ‘*‘Verification”* (Fig. 2, middle) experiments, BATEX configurations incorporating a HAM-like spring (H0, **<u>H</u>**R) yielded 6-7% metabolic cost reduction compared to not wearing an exo. Thus, simply placing biarticular springs bilaterally across the posterior thighs - even with a fixed stiffness (650 and 1100 N/m), is an effective across-users ‘generalizable’ approach that can universally reduce metabolic effort.

#### Selecting the per-participant ‘Best’ configuration (*Fig. 2*, Best)

Our “*Verification*” experiments (Fig. 2, right) point to the potential for additional advantages from uniquely configuring BATEX for each individual. In the average of two sets of experiments, there is a more pronounced metabolic reduction in the ’Best’ condition of approximately 9% compared to the fixed parameter condition (e.g., H0 with about 7% metabolic reduction), demonstrating the complementary benefits of BATEX’s spatial energy transfer and individualization effects. These findings highlight intersubject variability, emphasizing the need for personalized exosuit design. Additionally, inter-session variability may influence the consistency of improvements, underscoring the importance of advanced optimization techniques and training to refine parameter adjustments. Future research should explore the impact of training effects on longterm performance, as well as the integration of more autonomous Human-in-the-Loop Optimization [8] and the influence of human factors such as gender, mental states (e.g., confidence, trust, or stress), and other personal attributes in the interplay between the human and exosuit [57]. Addressing these aspects will be crucial for maximizing the exosuit’s effectiveness and user adaptability.

### 4.6 Muscle behavior adaptation explains response to assistance

Analyzing the impacts of the exosuit torque and mechanical power on users’ lower-limb joint kinematics, mechanics, and muscle activity highlights how compliant biarticular mechanisms enhance human walking economy with unpowered wearable systems. In this context, ‘morphological computation’ refers to the device reducing user effort (resulting from muscle activation) without requiring actuators, sensors, or external computational control. Any reduction in muscle activation further suggests a decrease in user control effort, achieved without additional computational cost.

The reduced RF EMG peak in the early swing phase (using RF springs) supports our hypothesis about the potential assistance of this biarticular spring for leg swinging. As discussed before, this supports faster swing leg protraction. A small (nonsignificant) increase in the HAM activation with these configurations (e.g., 0L or HL) could be an indirect effect of faster propulsion of the swing leg with the help of the RF spring. A more bent hip that can facilitate walking faster [58] is an advantage of using RF springs in BATEX. Using such a hip strategy to design and control exoskeletons has also shown long-term advantages in elderly adults’ gait [59].

Finally, the correlation between different muscle activations and the metabolic cost in Fig. 5 nicely supports the hypothesized contribution of the biarticular springs with BATEX. Our analysis clearly shows that the two biarticular muscles, rectus femoris (RF) and hamstrings (HAM), are the primary contributors to changes in metabolic cost, aligning with the design principles of BATEX. The next most important muscle is the next biarticular muscle GAS. The changes in GLM (as the hip extensor) activation can slightly improve our prediction of metabolic cost variations (see Fig. 5). This correlation measure is consistent with well-established physiological principles, where muscle activation typically leads to an increase in metabolic energy expenditure [48, 49, 60]. Enforcing positive coefficients in the regression model aligns with this principle, ensuring that the analysis remains physiologically realistic. The inclusion of all six monitored muscles was essential to capture the full scope of the relationship between muscle activity and metabolic cost. Additionally, the use of adjusted-*R*^2^ ensured a robust model fit without artificial improvements from non-contributory variables.

The design of the BATEX was inspired by template-based modeling [61] and the Locomotor Subfunctions (LSF) concept [62], showing the potential advantages of biarticular arrangements [19, 18] for supporting Swing and Balance LSF. The *Main Experiment* showed that adding either HAM or RF spring with low stiffness to the exo (H0 or 0R) could have significant effects on the metabolic cost. These two configurations, with only one engaged muscle per leg (either HAM or RF), could reduce metabolic costs in 7 and 8 subjects, for HAM and RF, respectively, even though their assisting principles are totally different. Among seven subjects who showed a similar decrease for both H0 and 0R configurations, all except one subject could benefit from **<u>H</u>**R (with both HAM and RF springs). This means that if subjects can benefit from HAM and RF biarticular muscles separately, with a chance of about 86% (6 out of 7), they can also experience metabolic reduction when using both artificial springs. Therefore, different assistance mechanisms of HAM and RF springs can be combined and support subjects with **<u>H</u>**R configuration. Although the metabolic reduction with this configuration was not significant in the *Main Experiments*, it could significantly reduce metabolic cost by more than 6% in *Verification* and more than 5% on average across two experiments. These results indicate that, compared to the single-spring configuration, the two-spring arrangement may present greater challenges for the user, potentially requiring a longer adaptation period to effectively utilize this assistance. This means that subjects could learn faster to take advantage of one of the temporal (by H0) or spatial (by 0R) energy transfers while benefiting from both (e.g., in **<u>H</u>**R) might need longer training time.

### 4.7 Outlook: Bioinspired Adaptive Body

This study presented a biologically inspired approach to human movement assistance by leveraging an unpowered system (i.e., *passive* BATEX) with key biomechanical features – specifically compliance and biarticularity– to assist and augment human locomotion. This framework opens new perspectives on movement assistance by assessing the extent of mechanical properties (i.e., morphological foundation) required and the role of muscle engagement in enabling adaptability across various movement tasks. As a next step to apply morphological computation to develop more adaptable assistive devices, we recently introduced the *active* BATEX device [41], featuring a bioinspired controller that underscores the importance of aligning mechanical responses between humans and robotic systems for smooth interaction. However, transitioning from passive to fully active systems requires energy, sensors, and sophisticated controllers. An alternative is to search for *adjustablity as needed* e.g., with quasi-passive systems [63, 64]. We envision to achieve this by enhancing BATEX’s adaptable body morphology. One potential parameter for tuning that can significantly affect GRF direction control for postural balance or influence leg swinging is the moment-arm ratio between different joints [19, 18]. In addition to a morphological foundation comprised of compliant inter-joint couplings (i.e., passive BATEX), engineers can integrate tunable materials [65] and a sensory feedback network (digital reflex pathways, [66, 67]) and then extend and fine-tune controllers based on gait template dynamics [42]. At the highest level, energy transfer among muscles, tendons, and compliant assistive elements needs to be optimized e.g., by Human-in-the-Loop optimization (HILO) [43].

Following our approach, two design and control dimensions emerge for assistive technologies: (1) tuning of passive mechanical features of the exoskeletal design and (2) calibrating the cooperativity between passive and active elements to shape and control user movement. True cooperation moves beyond mere assistance, creating new solutions for locomotion. In the biological body, each muscle-tendon complex is defined by the ratios between muscle-to-tendon length and muscle-to-tendon crosssectional area [68]. Developing bioinspired adaptable and customizable mechanical frameworks such as electric-pneumatic actuators (EPA) [69] offers a comparable solution for technical locomotor systems with minimal control effort while preserving muscle-like behavior [70]. However, the rules for individually adjusting each element’s design features, as in biological muscles, remain unclear. Identifying such self-tuning mechanisms *combining morphological and neural learning* could be an important step towards user-specific designs of assistive systems. Advanced multi-objective HILO techniques [71, 72] may refine the parameters of quasi-passive assistive systems to enhance locomotion metabolic efficiency, performance or other relevant measures such as agility and robustness. Such advanced optimization methods for daily activities (e.g., [73]) could propel human-robot interaction towards truly assistive and augmentative systems. This approach introduces intelligent biomechanics at a foundational level, integrating adaptability through material science [74] and potentially incorporating reflex-based topologies, facilitating multi-joint connections via structural elements (e.g., tunable muscles) and control systems (e.g., neural connectivity) [75].

## 5 Declarations

### 5.1 Ethics approval and consent to participate

This study was approved by the Ethical Committee of the Technical University of Darmstadt and was carried out based on the guidelines of the Declaration of Helsinki.

### 5.2 Consent for publication

All participants voluntarily provided written informed consent before participation in this study.

### 5.3 Availability of data and materials

The processed dataset of the here presented experimental study is publicly available in [76].

### 5.6 Author contributions

V.F. managed data curation, analysis, and interpretation, prepared the graphs, and contributed to writing the paper. A. A developed the hardware setup of the BATEX and contributed to conducting the experiments, data analysis, and developed figures. A. D. conducted the experiments and contributed to data collecting and processing. D. H contributed to experimental measurement and data collection. A. S. is the head of the Lauflabor Locomotion lab where the study is conducted and contributed to supervising research, discussions, data interpretation, writing the paper and funding acquisition. G. S. S. contributed to supervising the study, data analyses and interpretation, writing, review & editing of the manuscript. M. A. S. is the corresponding author of the article, responsible for the conceptualization and design of experiments, analyses and interpretation of data, and writing, review & editing of the manuscript, and funding acquisition.

## Acknowledgements

We thank members of the Lauflabor Locomotion lab for their assistance with data collection and constructive discussions.

## 6 Supplementary Information

### 6.1 Supplementary Text

This part includes supplementary figures and a table which complement the content of main text.

**Fig. S1.**
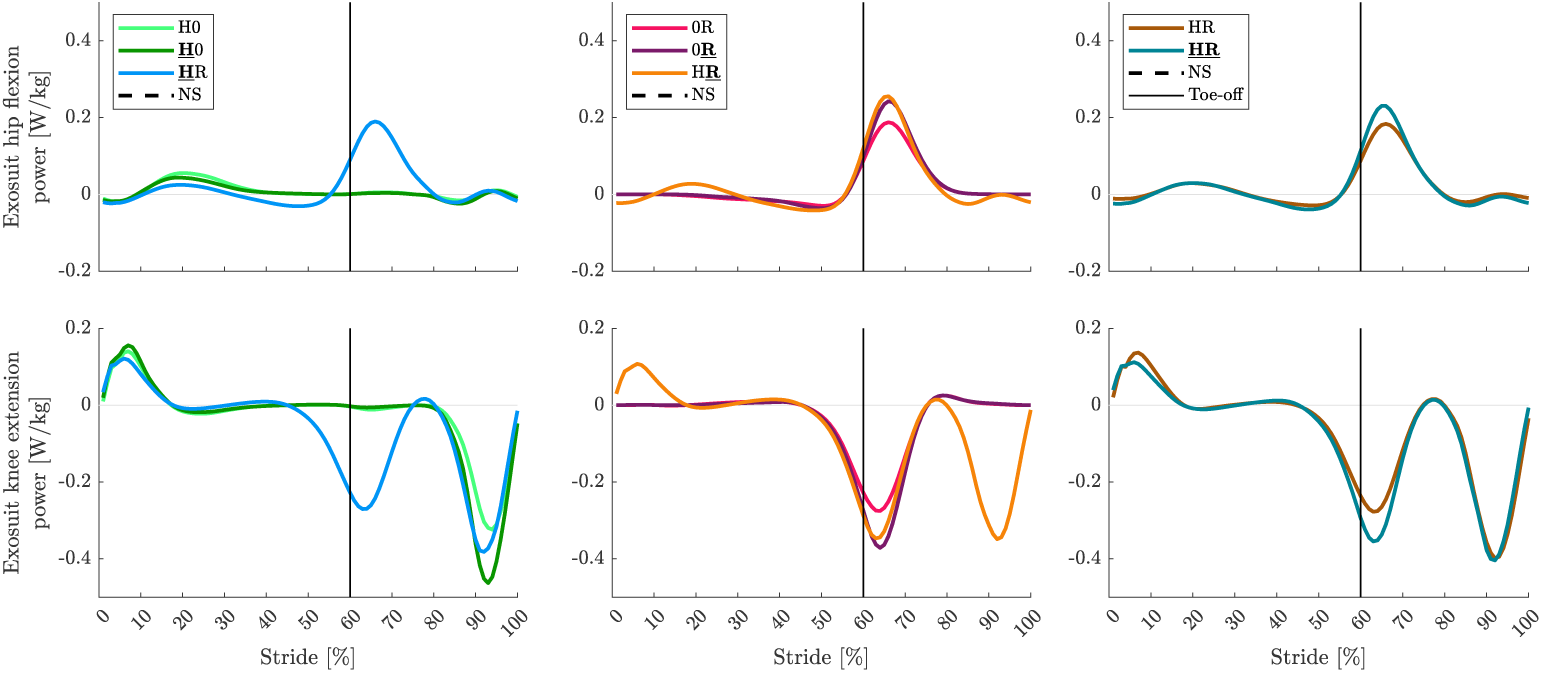
Exosuit power: Average exosuit power (normalized to body mass) at the hip, and knee joints of nine participants in one stride, comparing different BATEX springs’ combinations effects. The first column illustrates configurations with higher stiffness in HAM springs compared to RF springs. The second column depicts configurations with greater RF stiffness compared to HAM. The last column showcases configurations with equal stiffness for both HAM and RF. The vertical black line shows the toe-off moment. The stride starts at the right leg touch-down and ends at the touch-down of the same leg.

**Fig. S2.**
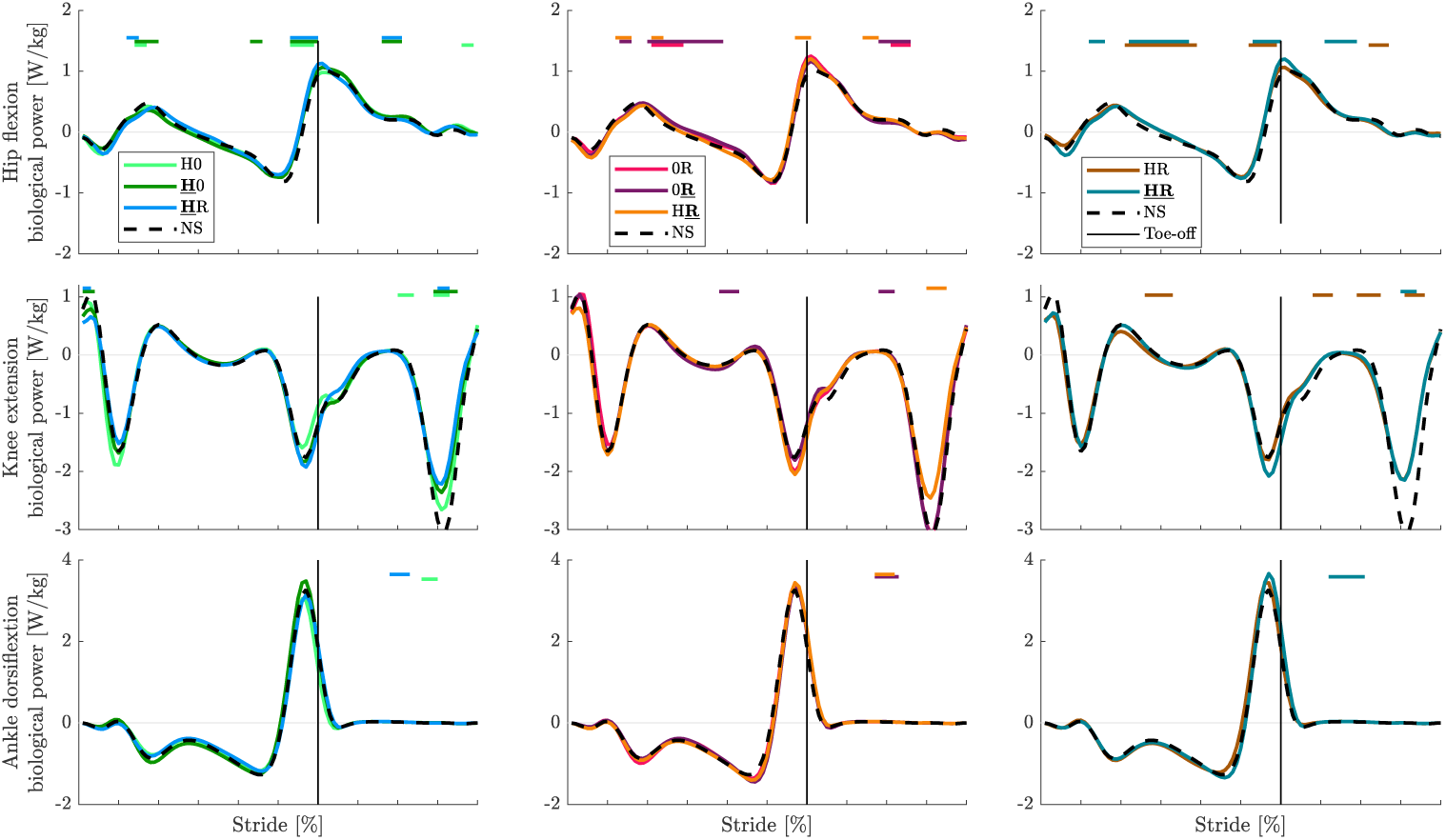
Biological joint power: Average biological join power (normalized to body mass) at the hip, knee, and ankle joints of nine participants in one stride comparing different BATEX springs’ combinations effects. The first column illustrates configurations with higher stiffness in HAM springs compared to RF springs. The second column depicts configurations with greater RF stiffness compared to HAM. The last column showcases configurations with equal stiffness for both HAM and RF. The NS case is represented in all configurations with a black dashed line, as a baseline for comparison. The vertical black line shows the toe-off moment. The stride starts at the right leg touch-down and ends at the touch-down of the same leg. Statistically significant differences (*p <* 0.05) between NS and other exosuit configurations are highlighted with horizontal lines above each figure.

**Fig. S3.**
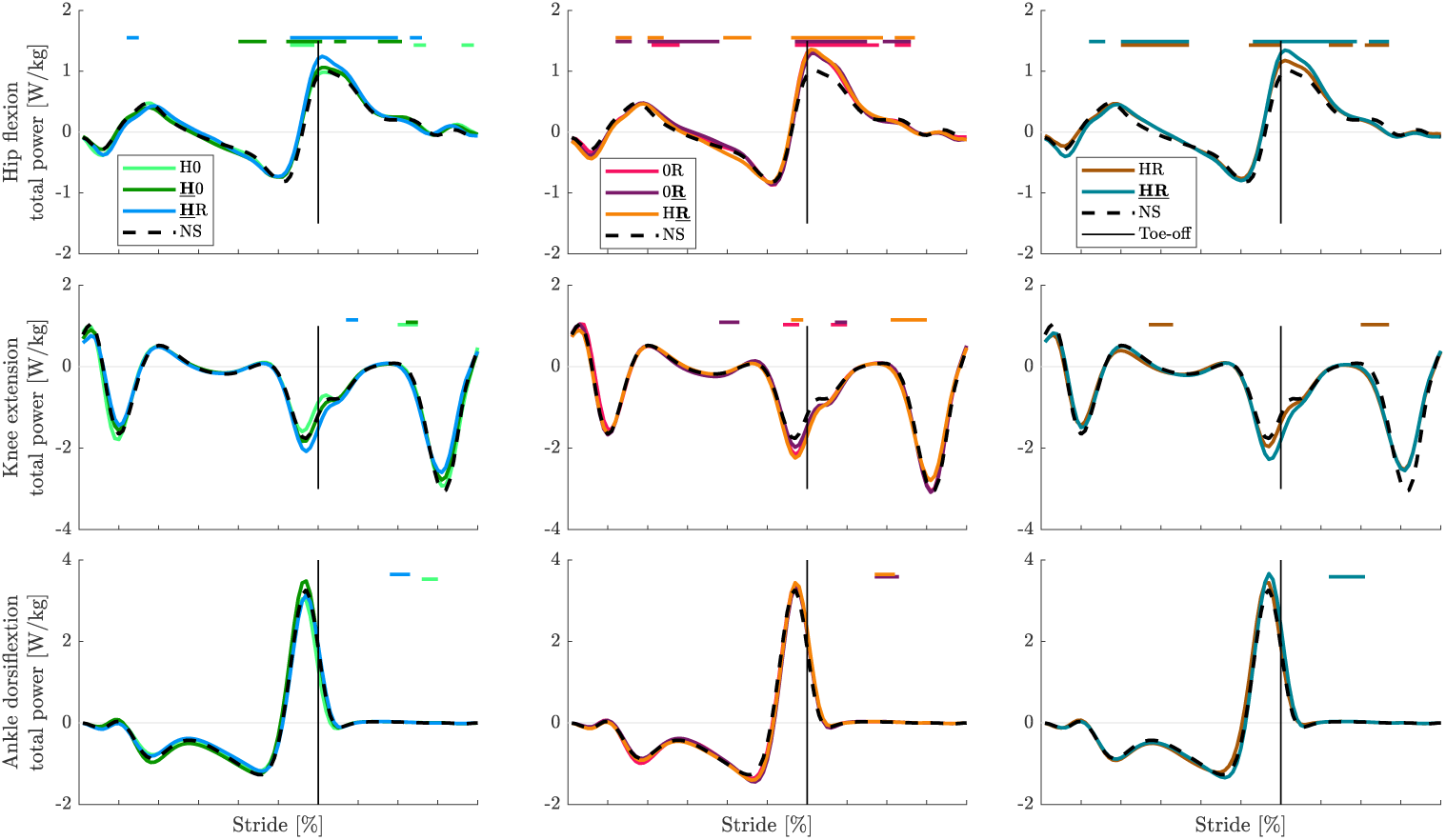
Total joint power: Average total joint power (normalized to body mass) at the hip, knee, and ankle joints of nine participants in one stride comparing different BATEX springs’ combinations effects. The first column illustrates configurations with higher stiffness in HAM springs compared to RF springs. The second column depicts configurations with greater RF stiffness compared to HAM. The last column showcases configurations with equal stiffness for both HAM and RF. The NS case is represented in all configurations with a black dashed line, as a baseline for comparison. The vertical black line shows the toe-off moment. The stride starts at the right leg touch-down and ends at the touch-down of the same leg. Statistically significant differences (*p <* 0.05) between NS and other exosuit configurations are highlighted with horizontal lines above each figure.

**Fig. S4.**
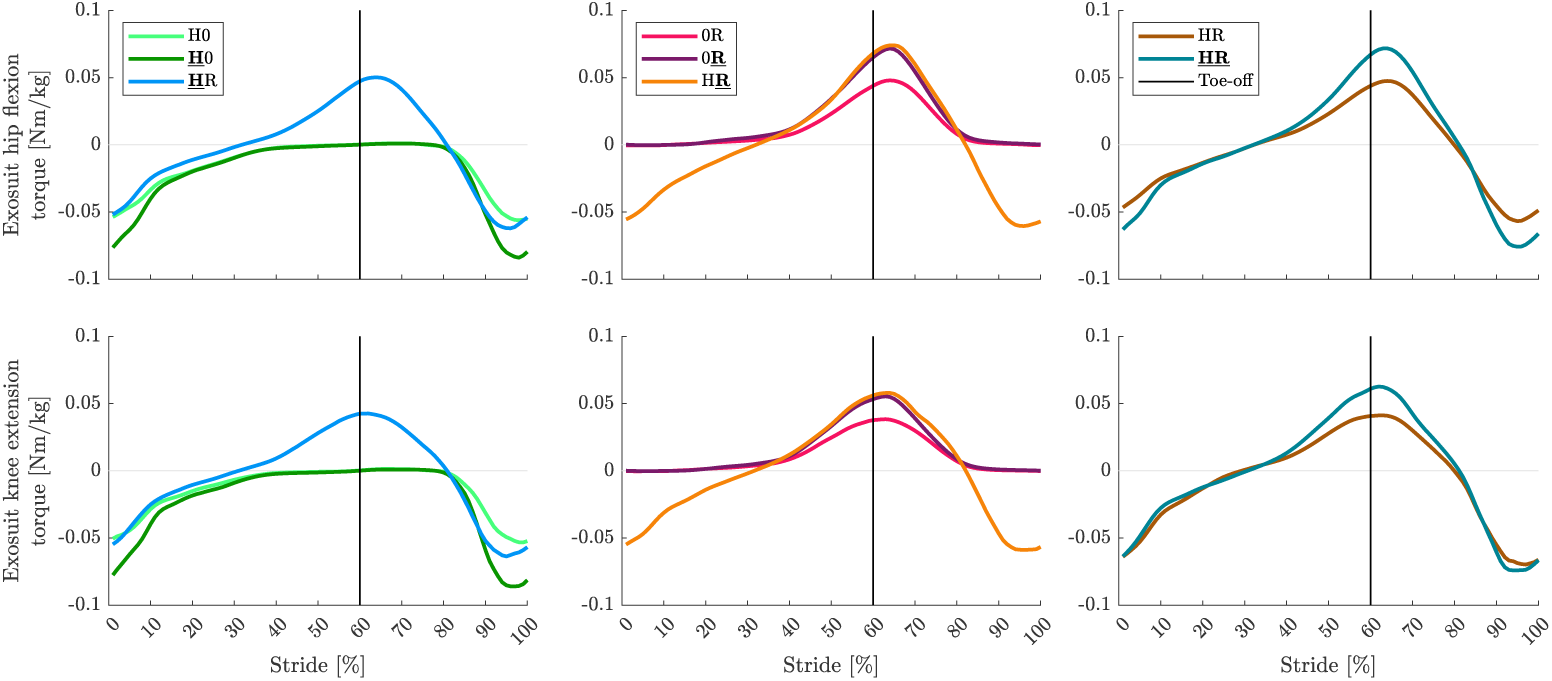
Exosuit torque: Average exosuit torque (normalized to body mass) at the hip, and knee joints of nine participants in one stride, comparing different BATEX springs’ combinations effects. The first column illustrates configurations with higher stiffness in HAM springs compared to RF springs. The second column depicts configurations with greater RF stiffness compared to HAM. The last column showcases configurations with equal stiffness for both HAM and RF. The vertical black line shows the toe-off moment. The stride starts at the right leg touch-down and ends at the touch-down of the same leg.

**Fig. S5.**
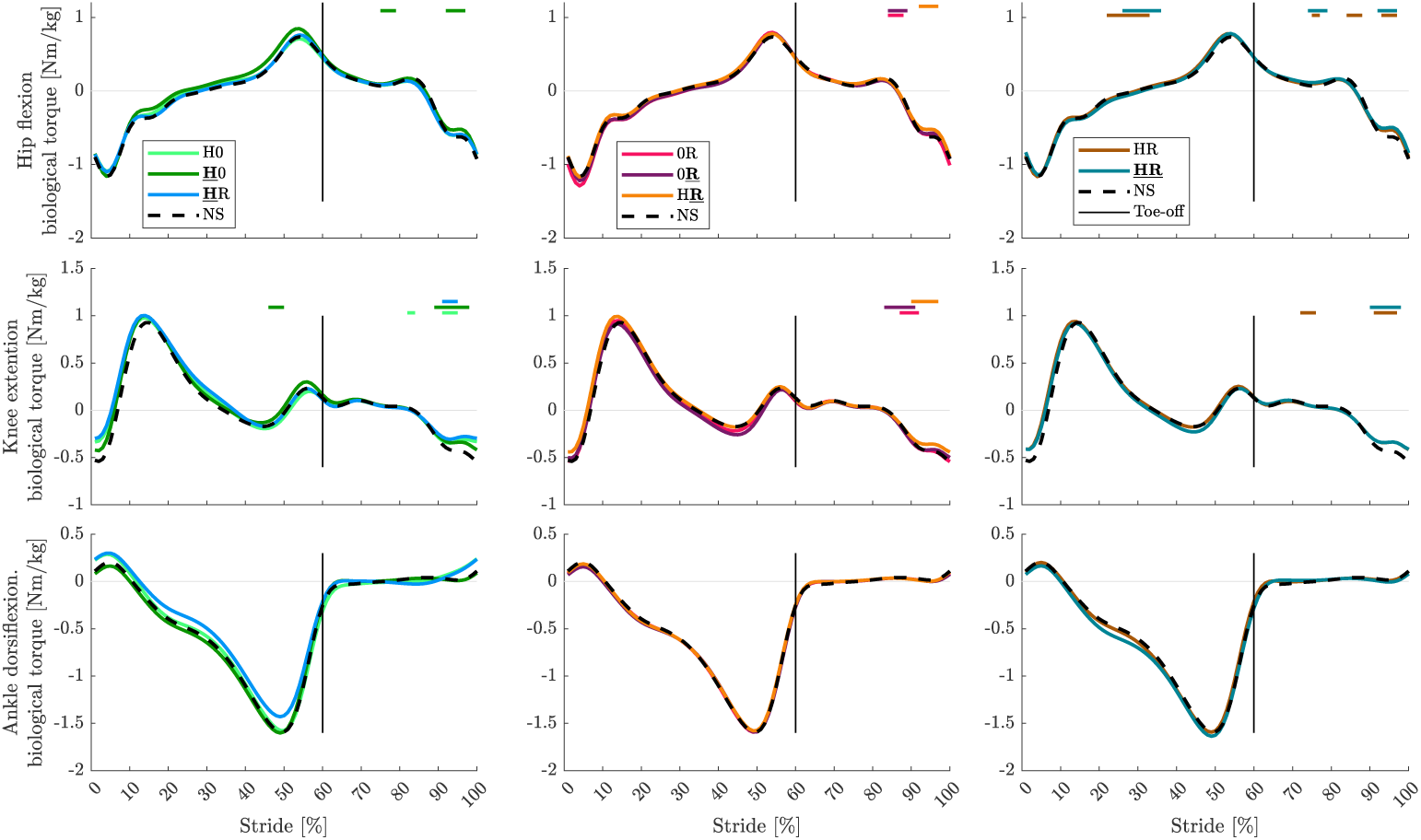
Biological joint torque: Average biological joint torque (normalized to body mass) at the hip, knee, and ankle joints of nine participants in one stride comparing different BATEX springs’ combinations effects. The first column illustrates configurations with higher stiffness in HAM springs compared to RF springs. The second column depicts configurations with greater RF stiffness compared to HAM. The last column showcases configurations with equal stiffness for both HAM and RF. The NS case is represented in all configurations with a black dashed line as a baseline for comparison. The vertical black line shows the toe-off moment. The stride starts at the right leg touch-down and ends at the touch-down of the same leg. Statistically significant differences (*p <* 0.05) between NS and other exosuit configurations are highlighted with horizontal lines above each figure.

**Fig. S6.**
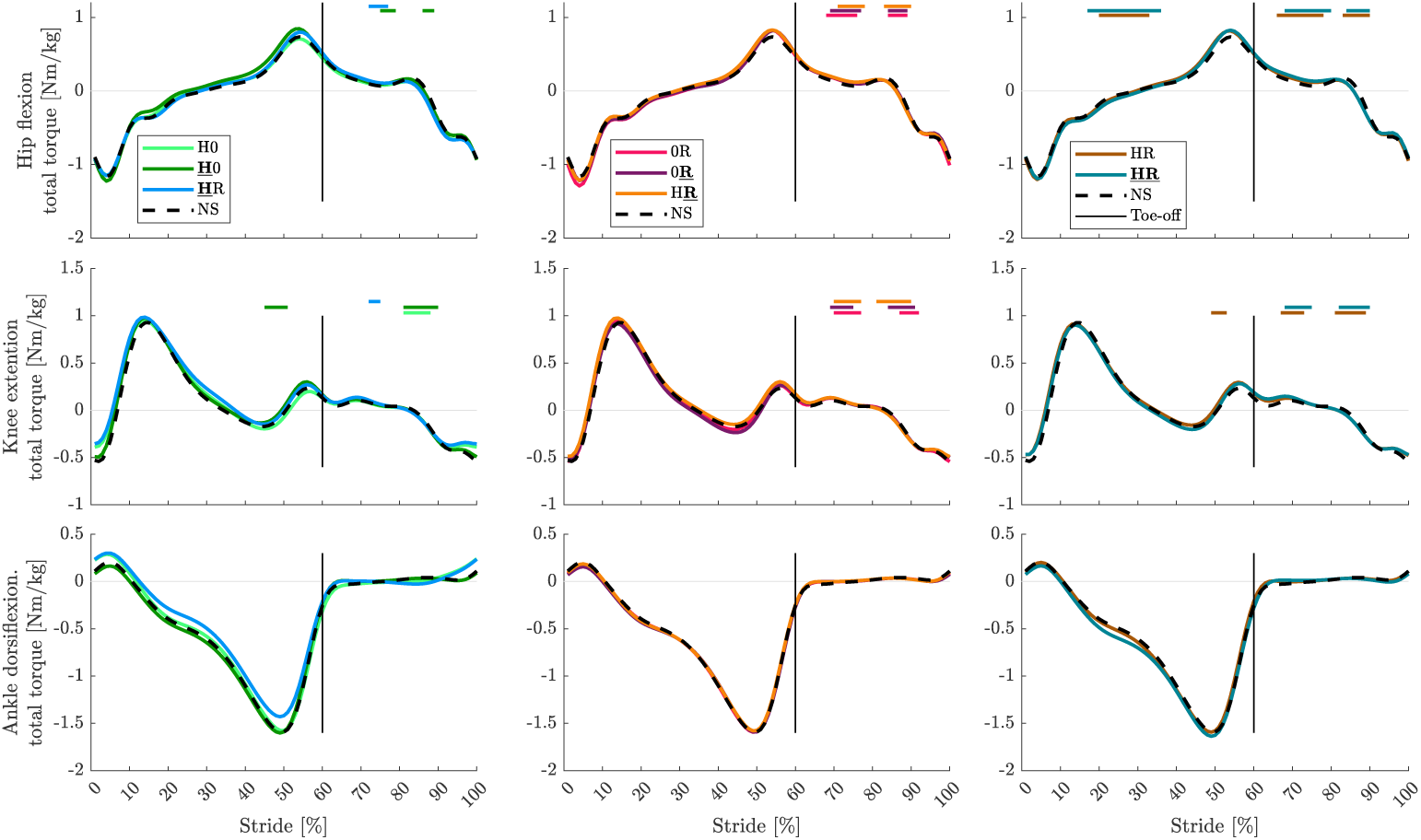
Total joint torque: Average total joint torque (normalized to body mass) at the hip, knee, and ankle joints of nine participants in one stride comparing different BATEX springs’ combinations effects. The first column illustrates configurations with higher stiffness in HAM springs compared to RF springs. The second column depicts configurations with greater RF stiffness compared to HAM. The last column showcases configurations with equal stiffness for both HAM and RF. The NS case is represented in all configurations with a black dashed line, as a baseline for comparison. The vertical black line shows the toe-off moment. The stride starts at the right leg touch-down and ends at the touch-down of the same leg. Statistically significant differences (*p <* 0.05) between NS and other exosuit configurations are highlighted with horizontal lines above each figure.

**Fig. S7.**
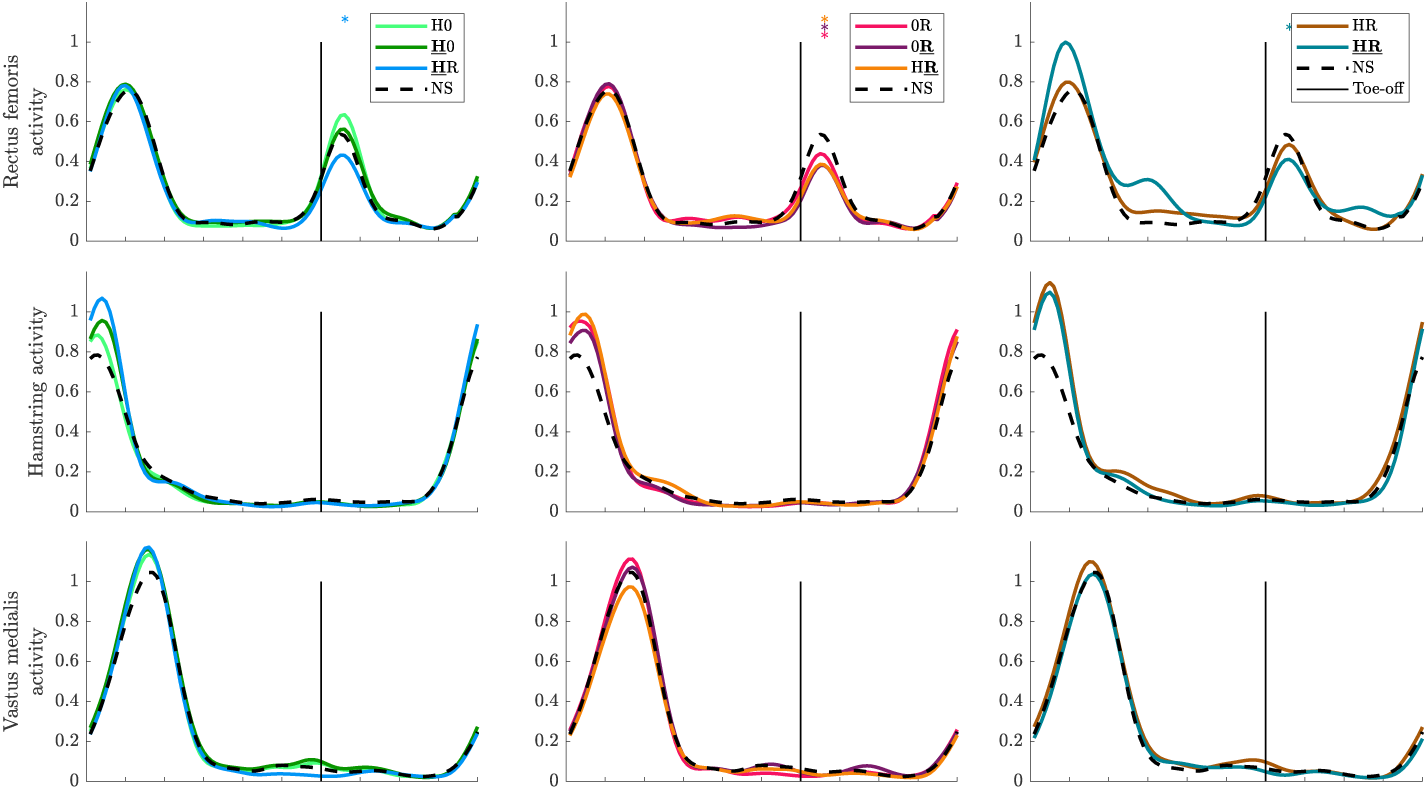
Lower limb proximal muscle activity: Average Muscle activities in the rectus femoris (RF), hamstring (HAM), and vastus medialis (VAS) of nine participants in one stride comparing different BATEX springs’ combinations effects. Muscle activities are normalized to the average peak of NS among participants. The first column illustrates configurations with higher stiffness in HAM springs compared to RF springs. The second column depicts configurations with greater RF stiffness compared to HAM. The last column showcases configurations with equal stiffness for both HAM and RF. The NS case is represented in all configurations with a black dashed line, as a baseline for comparison. The vertical black line shows the toe-off moment. The stride starts at the right leg touchdown and ends at the touch-down of the same leg. Significant changes in EMG peak (*p <* 0.05) are marked with an asterisk (*).

**Fig. S8.**
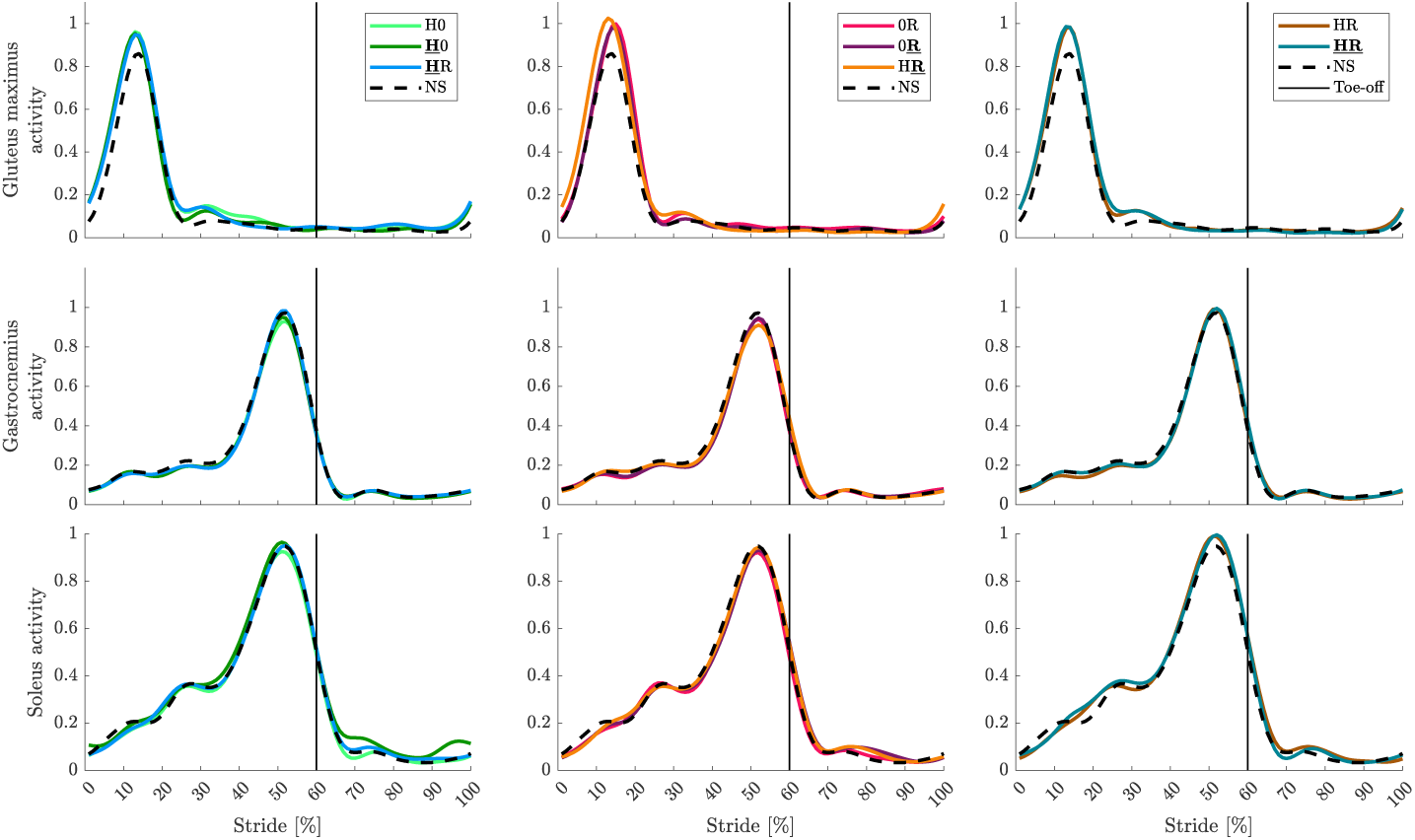
Lower limb distal muscle activity: Average Muscle activities in gluteus maximus (GLM), gastrocnemius (GAS), and soleus (SOL) of nine participants in one stride comparing different BATEX springs’ combinations effects. Muscle activities are normalized to the average peak of NS among participants. The first column illustrates configurations with higher stiffness in HAM springs compared to RF springs. The second column depicts configurations with greater RF stiffness compared to HAM. The last column showcases configurations with equal stiffness for both HAM and RF. The NS case is represented in all configurations with a black dashed line, as a baseline for comparison. The vertical black line shows the toe-off moment. The stride starts at the right leg touch-down and ends at the touch-down of the same leg. Significant changes in EMG peak (*p <* 0.05) are marked with an asterisk (*).

**Fig. S9.**
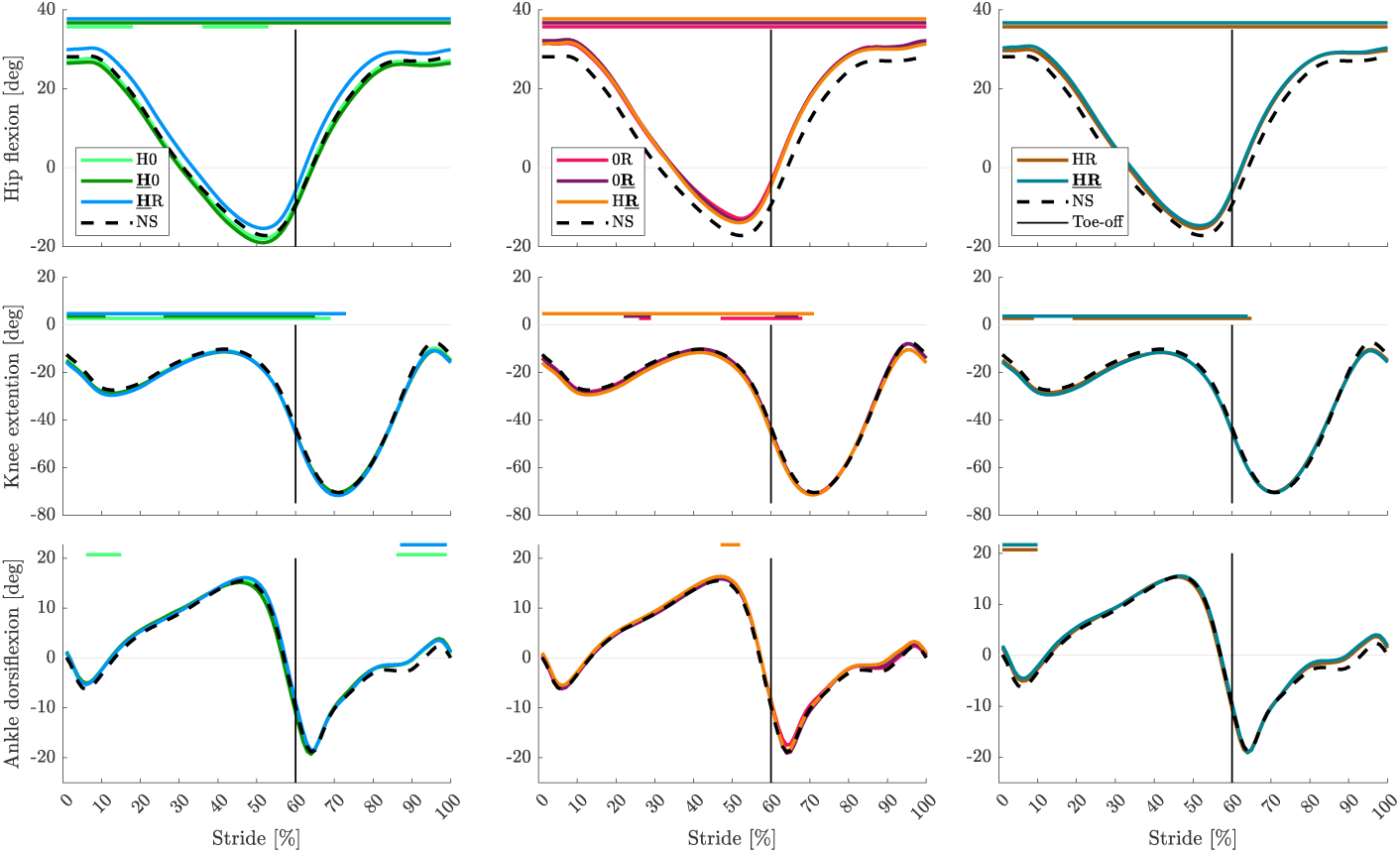
Joint angle: Average total joint angle at the hip, knee, and ankle joints of nine participants in one stride comparing different BATEX springs’ combinations effects. The first column illustrates configurations with higher stiffness in HAM springs compared to RF springs. The second column depicts configurations with greater RF stiffness compared to HAM. The last column showcases configurations with equal stiffness for both HAM and RF. The NS case is represented in all configurations with a black dashed line, as a baseline for comparison. The vertical black line shows the toe-off moment. The stride starts at the right leg touch-down and ends at the touch-down of the same leg. Statistically significant differences (*p <* 0.05) between NS and other exosuit configurations are highlighted with horizontal lines above each figure.

**Fig. S10.**
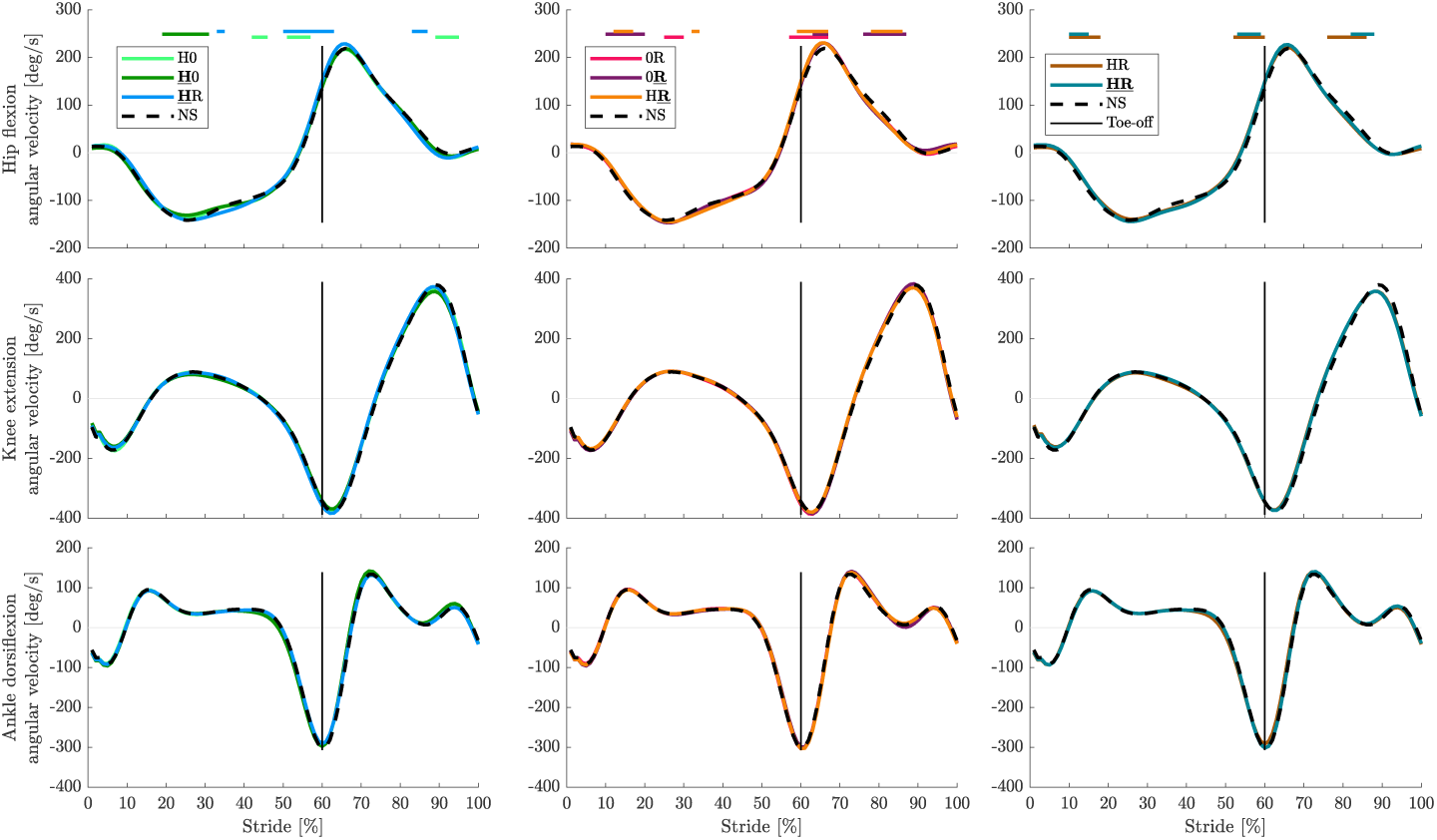
Joint angular velocity: Average total joint angular velocity at the hip, knee, and ankle joints of nine participants in one stride comparing different BATEX springs’ combinations effects. The first column illustrates configurations with higher stiffness in HAM springs compared to RF springs. The second column depicts configurations with greater RF stiffness compared to HAM. The last column showcases configurations with equal stiffness for both HAM and RF. The NS case is represented in all configurations with a black dashed line, as a baseline for comparison. The vertical black line shows the toe-off moment. The stride starts at the right leg touch-down and ends at the touch-down of the same leg. Statistically significant differences (*p <* 0.05) between NS and other exosuit configurations are highlighted with horizontal lines above each figure.

**Table S1.** Lever arms specifications: Maximum and minimum lever arm lengths (mean ± standard deviation) for the hip and knee joints, measured for the RF and HAM springs.

| Joint | Lever arm (max) | Lever arm (min) |
| --- | --- | --- |
| Hip (RF) | $14.7 \pm 2.1$ | $12.35 \pm 1.6$ |
| Hip (HAM) | $15.9 \pm 2.4$ | $13.3 \pm 1.25$ |
| Knee (RF) | $15.3 \pm 2.35$ | $10.4 \pm 1.5$ |
| Knee (HAM) | $13.6 \pm 3$ | $11.7 \pm 2.05$ |

An appendix contains supplementary information that is not an essential part of the text itself but which may be helpful in providing a more comprehensive understanding of the research problem or it is information that is too cumbersome to be included in the body of the paper.

